# Prediction of mosquito vector abundance for three species in the *Anopheles gambiae* complex

**DOI:** 10.1101/2025.03.04.641552

**Authors:** Geoffrey R. Hosack, Maud El-Hachem, Adrien Ickowicz, Nicholas J. Beeton, Andrew Wilkins, Keith R. Hayes, Samuel S. C. Rund, Sarah A. Kelly, Mary Ann McDowell

## Abstract

**Background:** The dynamics of vector borne disease transmission depend on the abundances of vectors. The dominant malaria vector species complex of *Anopheles gambiae sensu lato* is a target of vector control strategies designed to reduce and eliminate malaria transmission. The three most widely distributed dominant malaria vectors within the species complex are *An. arabiensis, An. coluzzii* and *An. gambiae sensu stricto*.

**Methods:** Previous studies across the extent of the species complex range have been restricted to annual predictions of species occurrence or relative abundance. This study incorporated relative abundance data at the species level and abundance data at the species complex level to estimate and predict daily equilibrium vector abundances of each species. Statistical models with interpretable parameters were used to gain insight into how factors such as meteorological conditions, insecticide treated net use, and human density affect the spatial and temporal predictions. The statistical models were used to predict daily equilibrium vector abundance that is an important factor in indices of malaria transmission such as the basic reproduction number and vectorial capacity.

**Results:** Meteorological factors, such as precipitation and relative humidity, and human factors, such as population density and insecticide treated nets, were important predictors of these three malaria vector species in Africa. Insecticide treated net use was negatively associated with per capita vector abundance of the *An. gambiae* species complex and interacted with year to suggest an a negative effect on the ratio of *An. gambiae s.s*. to *An. arabiensis* at the end of the prediction period that extended from 2002 to 2020. A predicted increasing interannual trend for *An. coluzzii* was potentially caused by changing species identification methods.

**Conclusion:** The predicted equilibrium abundances of the three species showed potentially high levels of geographical overlap, niche overlap, and evidence for stable coexistence despite substantial interspecific competition. Improving collection of longitudinal species abundance data across the spatial range of the *An. gambiae* species complex will facilitate future explorations of causal hypotheses that relate vector abundance to control measures, malaria interventions and meteorological conditions.

## Background

The reduction of mosquito vector abundance and the number of infectious bites on humans through vector control measures is an effective means to lower or eliminate human malaria [1–3]. Shifts in species abundances of mosquito vectors induce concomitant changes in behaviour and susceptibility to vector control that require adjustment of available intervention strategies such as insecticide treated nets, indoor residual spraying, larval source management, house screening, baited traps or novel techniques such as genetically modified vector suppression and replacement strategies [4–6]. The evaluation of conventional and potential novel vector control strategies on disease transmission dynamics thus benefits from prediction and estimation of vector abundances [7, 8]. Mathematical models of long term or equilibrium vector abundance, for example, have long served as useful guides for vector control and malaria interventions [9]. Equilibrium vector abundance is the theoretical population size that would be attained if birth rates matched death rates at a given location. Equilibrium abundance is a useful concept allowing for the possibility that the true or actual abundance of a species may diverge from carrying capacity because of local environmental or anthropogenic perturbations, dispersal, source-sink meta-population dynamics or through interspecific competition [10–12]. For example, vector populations fluctuate around the equilibrium in response to seasonal changes in birth and death rates, and a range of other factors.

Applications with a time-varying basic reproduction number (ℛ _0_) that depends on the ratio of vector abundance to human hosts, and derived indices such as time-varying vectorial capacity, are examples of an equilibrium based analysis. ℛ_0_ is defined at the disease free equilibrium composed of completely susceptible vector and human populations in the standard Ross-Macdonald model [1, 2, 13] and related mathematical models of malaria transmission by anophelines [14]. The canonical basic reproduction number ℛ_0_ corresponds to the number of malaria infections that arise from a single infectious case introduced into a completely susceptible population at the disease free equilibrium [15]. Macdonald acknowledges that the changing vector abundance and transmission parameters that define ℛ_0_ suggest that “… the degrees of equilibrium seen in nature should make a continuous series …” (p. 34 in [2]). The equilibrium index ℛ_0_, for example, can be modelled as a continuous function of temperature and precipitation mediated by the average vector population at a climatically determined equilibrium [16]. The influential Garki model [9, 17, 18] accounts for seasonal malaria transmission in Nigeria via an interpolated function of time-varying vector population size that determines the vector population’s contribution to ℛ_0_ through the vectorial capacity [19]. Recent statistical analysis of the Garki data from the 1970s suggests that the equilibrium vector population size of *An. gambiae s.l*. defined by an environmentally driven carrying capacity approximately tracks weekly precipitation [20]. An analysis of seasonal malaria transmission in Burkina Faso suggests that the average vector population size of *An. gambiae* is proportional to weekly precipitation [21].

Despite the important contribution of the vector-to-human host ratio to the standard indices of malaria transmission risk such as the basic reproduction number ℛ_0_ and vectorial capacity, abundance data for mosquito vector species are rarely collected in Africa [22]. In particular, the species complex *Anopheles gambiae sensu lato* (*s.l*.) is an important vector of malaria in Africa [23]. *An. gambiae s.l*. is estimated to have caused at least half of the 241 million malaria deaths of children under five years old in sub-Saharan Africa between 3000 BCE and 2000 CE [6]. The three primary malaria vector member or sibling species of the *An. gambiae* complex [6] that have the broadest geographical distribution [24] are *An. arabiensis, An. coluzzii* and *An. gambiae sensu stricto* (*s.s*.). However, longitudinal abundance data for species in the *An. gambiae* complex are rare. Species identification in field collections is often limited because the member species are not morphologically distinguishable, and so molecular identification from subsamples of field collections is required. Longitudinal count data for *An. arabiensis, An. coluzzii* and *An. gambiae s.s*. are therefore uncommon, and most predictive studies at the continental scale rely only on species occurrence data [24–27].

Predictions of the probability of species occurrence from species distribution models link the distributions of species in the *An. gambiae* complex to environmental covariates such as annual indices of meteorological conditions and climate [24–26]. Alternatively, climatological covariates may be used to predict relative abundance or species composition [28] based on species occurrence data [27, 29]. Sinka et al. [27] suggest that the consideration of relative abundance may defray some of the difficulties associated with the estimation of ‘true’ abundance that is spatially and temporally variable, and use species occurrence data to predict that the relative abundance of *An. arabiensis* versus the aggregate of the two species *An. coluzzii* and *An. gambiae s.s*. increases with insecticide treated net intervention. Tene Fossog et al. [29] use relative species frequencies of *An. coluzzii* and *An. Gambiae s.s*. to predict relative probabilities of occurrence in western Africa based on climatic and geographical covariates with species specific responses determined by elevation and distance from coastline. Pombi et al. [30] find high niche overlap among *An. arabiensis, An. coluzzii* and *An. gambiae s.s*. in certain regions based on an index that is a function of relative abundance data [31]. Niche overlap is sometimes suggested as an index of the strength of interspecific competition [32], although this correlative index based on the relative abundance of species may diverge from the underlying competitive relationships among species [31]. Competition among different species of mosquito larvae can lead to the competitive displacement or reduction of one species by another and so affect strategies for control of disease vectors [33, 34].

However, malaria transmission dynamics are driven by the abundance of mosquito species and not just relative abundance or the probability of species occurrence. This study therefore sought to estimate spatiotemporal abundance of *An. arabiensis, An. coluzzii* and *An. gambiae s.s*. by collating data both on relative abundance at the species level (identified genotypically) and longitudinal population counts at the species complex level (identified morphologically) across the geographic range of *An. gambiae s.l*. The difficulty in acquiring such data should not be underestimated or understated. Regular vector surveillance is typically conducted to support operational activities, which in the African entomological context is overwhelmingly in support of malaria intervention through insecticide treated bednet (ITN) deployment or indoor residual spray (IRS) activity. Indeed, much of the regular surveillance is focused on determining local species present and their localised behaviours (e.g., measures of anthropophily, endophily, and biting time) to determine appropriate vector interventions [35]. Unlike some regions that employ weekly or daily surveillance to identify mosquito control action thresholds for vector suppression operations such as insecticidal fogging, the operational timescales for African ITN distributions or IRS operations are not based on the weekly or monthly status of mosquito species abundance. Hence, such data are less likely to be collected. This fact is highlighted in a recent global meta-analysis of mosquito control action thresholds [36], where none are listed from Africa. Consequently, although longitudinal surveillance of *Anopheles* mosquito populations is performed in Africa particularly for research purposes, most of the longitudinal studies investigated in this study either did not use genetic methods (e.g., polymerase chain reaction, PCR) to identify field collections to the species level or instead did not resolve changes in species diversity and relative abundance within a year. The ability to detect seasonal changes in species composition was therefore negated. Nonetheless, the species identified data collated by this study expand on previous efforts [24] by including finer scale temporal information on the sample collection time that thereby allows incorporation of seasonally varying environmental conditions into the analysis. Importantly, this data acquisition effort also currently represents the largest collation of publicly available longitudinal abundance data for the *An. gambiae* species complex. These data investigation and collation efforts both highlight and help bridge a significant gap in available data for African malaria mosquito biology.

Using these assembled data, the equilibrium abundances of *An. arabiensis, An. coluzzii* and *An. gambiae s.s*. were predicted across sub-Saharan Africa from 2002 to 2020. The species equilibrium abundances are not directly observable and were inferred from empirical observations as responding to known spatial and spatiotemporal covariates. Statistical relationships were estimated for established environmental and geographical factors, such as precipitation, temperature and human population density [16, 20, 21, 37], to enable spatiotemporal predictions of *An. gambiae s.l*. abundance. Previous relative abundance analyses for pairs of species in *An. gambiae s.l*. [27–29] were extended in this study to account for daily varying meteorological predictors to allow seasonally varying predictions of relative abundance for *An. arabiensis, An. coluzzii* and *An. gambiae s.s*. The analysis of relative abundance data available at the sibling species level was combined with the longitudinal abundance analysis at the species complex level to predict potential sibling species abundance across sub-Saharan Africa. The predicted equilibrium abundances of *An. arabiensis, An. coluzzii* and *An. gambiae s.s*. were then connected to the equilibrium of an underlying dynamic model that suggests the potential for strong interspecific competition but also stable coexistence among the three vector species.

## Methods

The study area is defined as the geographic range of *An. gambiae s.l*. [24, 38] in sub-Saharan Africa. The genotype data, abundance data, spatial covariates and spatiotemporal covariates are described below with further details provided in the supplementary R package **AgslPredict** for this paper [39]. The code for analyses and graphical results of relative abundance predictions, abundance predictions and competition estimates are also available in **AgslPredict**.

### Genotype and Abundance Datasets

The two sets of data, species level genotype data and species complex level abundance data, have been made publicly available at VectorBase.org and are each discussed in turn. The genotype data of the three species *An. arabiensis, An. coluzzii* and *An. Gambiae s.s*. were downloaded from VectorBase.org [40, 41] and spatially subsetted to the study area. Some samples of the *An. gambiae s.l*. species complex were identified by chromosomal form to species. Species recorded as “Anopheles gambiae chromosomal form Bamako” and “Anopheles gambiae chromosomal form Savanna” were assigned to “Anopheles gambiae sensu stricto”, and “Anopheles gambiae chromosomal form Mopti” was assigned to “Anopheles coluzzii” [42]. The three species retained for analysis were “Anopheles arabiensis”, “Anopheles coluzzii” and “Anopheles gambiae sensu stricto”. Samples for which information on date of collection was missing were filtered out of the dataset. Information on time of collection varied across the remaining samples but could include durations of a single day, month, year or multiple years. The start and end dates of each collection were extracted. Samples with collection dates reported only to the month were assigned start and end dates that corresponded to the start and end dates of the reported month. Samples with reported durations greater than 31 days were filtered out of the dataset. *An. coluzzii* and *An. gambiae s.s*. were not distinguished prior to 2001 [30] and samples with earlier initial dates were omitted. Samples were then identified by project ID, latitude and longitude, collection date and collection protocol to define the sample unit of analysis. The frequency counts of the three species identified for each unit sample were then recorded for use in analyses of species relative abundance (species composition) described in the next section. Sample units with shared collection dates and collection methods were aggregated to 5 km *×* 5 km grid cells across the spatial extent, which resulted in 1,740 records. The 5 km *×* 5 km grid cell resolution was chosen to capture the home range of a female mosquito since most mosquitoes disperse less than 5 km based on mark release recapture experiments [43] and for comparison with previous spatial analyses of genotyped data for *An. gambiae* species that also used 5 km *×* 5 km grid cell resolution [23, 24, 27, 29]. The collated records extended from 2001 to 2021 and are summarised in Table S1 and Figure S1.

The abundance data for *An. gambiae s.l*. were downloaded from VectorBase.org [40, 44] and spatially subsetted to the study area. Whereas the relative abundance data are typically derived from genotyped subsamples of field collections, the abundance data recorded from a field collection are most often counts of individuals morphologically identified to the level of the *An. gambiae* species complex. The observed counts of *An. gambiae s.l*. were treated as the response variable for the abundance analysis. Observation methods that could be categorised to specific collection methods were retained for analysis. The level of sampling effort was defined as the daily number of field collections by observation method in each grid cell. Records were then subsetted to samples identified as adult female mosquitoes. The available records extended from 1997 to 2021. Sites with long term longitudinal observations often also included long periods without observations. About 2% of the remaining samples had indeterminate sampling dates and were excluded from the analysis. Observations were aggregated to each grid cell by day and observation method, which resulted in 9,105 samples of abundance after excluding collection methods that targeted larvae or had low information content due to very few samples collected (less than ten records). The collated records are summarised in Table S2. These *An. gambiae s.l*. abundance records were sparsely distributed across sub-Saharan Africa (Figure S2).

### Covariate Datasets

Precipitation, temperature and humidity are meteorological predictors of *An. gambiae* distribution [24, 26, 45]. Spatiotemporal meteorological data for daily precipitation totals, daily average temperature and daily average relative humidity were obtained from the National Aeronautics and Space Administration (NASA) Langley Research Center (LaRC) Prediction of Worldwide Energy Resource (POWER) Project funded through the NASA Earth Science/Applied Science Program [46] using the R package **nasapower** [47, 48]. These data are provided at a resolution of 0.625° longitude by 0.5° latitude. The meteorological covariates of a meteorological grid cell were assigned to all those 5 km *×* 5 km grid cells with centroids that occurred within the corresponding meteorological grid cell. In the Supplementary Information, the meteorological data are visualised by annual average (Figures S3, S4, S5), quarterly average (Figures S6, S7, S8) and anomalies with respect to the average over the timespan 2002–2020 (Figures S9, S10, S11) used for the prediction period.

Distances to nearest coast, presence of freshwater, elevation, human population, insecticide treated net use, longitude and latitude are proposed spatial predictors of *An. gambiae* distribution [7, 24, 29, 37]. Spatial covariates were constructed using distances from 5 km *×* 5 km grid cell centroids to the nearest coast (Figure S12) using data provided by Natural Earth and obtained with the R packages rnaturalearth [49] and rnaturalearthdata [50]. Elevation data at 15 arcseconds resolution were obtained from the National Centers for Environmental Information, National Oceanic and Atmospheric Administration [51], and averaged to obtain mean elevation for each 5 km *×* 5km grid cell (Figure S13).

The presence of freshwater habitat was quantified by the probability of permanent fresh-water occurring in each 5 km *×* 5km grid cell, which was defined as follows. The minimum probability of intermittence for streams and rivers in each cell, which is derived from the predicted probability of flow cessation for at least 30 days per year on average [52], was first calculated for all stream and river reaches that occurred in a cell. The probability of permanent stream or river freshwater was then defined by the complementary probability, that is, one minus the minimum probability of intermittence. If a cell intersected with a lake or reservoir [LakeATLAS; 53], then the probability of freshwater was assigned a value of one. The probability of permanent freshwater is depicted in Figure S14.

Human population is often proposed as a predictor that is positively associated with the areal density of *An. gambiae* s.l. in rural environments [7, 24, 29, 37, 54]. However, this relationship breaks down in urban environments where malaria transmission is generally lower compared to rural and peri-urban environments [55, 56] and also more variable [57], noting that localised areas of high vector abundance near larval habitats can still occur in urban settings [55]. Total human population in each 5 km *×* 5km grid cell was therefore censored above 50,000 human inhabitants, which is the threshold commonly used to demarcate urban areas formed by contiguous areas with greater than 1,500 inhabitants per square kilometre [58]. The corresponding threshold for a censored human density covariate in each 5 km *×* 5km grid cell was censored at 2,000 inhabitants per square kilometre for consistency with the censored total human population covariate. The censored human population covariates therefore vary across rural and peri-urban 5 km by 5 km grid cells, and are assigned their threshold values in urban cells. Annual human population data at 30 arcseconds resolution were obtained from Landscan (Oak Ridge National Laboratory, https://landscan.ornl.gov/), and averaged to obtain total human population and mean human population density per square kilometere for each 5 km by 5 km grid cell (Figure S15).

Annual vector intervention data for ITN percent use by individuals and IRS percent coverage of households is available for the study area across the years 2000–2022 [59–61]. The intervention data were averaged to the 5 km *×* 5 km grid cells to derive spatiotemporal covariates of intervention effort. Caution is required for assessment of intervention efficacy on the basis of observational studies [62]. For example, intervention effort may increase for locations and times with high vector densities. A spurious positive association could then be induced between vector populations and intervention effort that thereby masks an expected negative causal impact of the intervention. Such an outcome could even be likely where interventions are selectively applied, but possibly less likely or severe for broadly applied interventions with deployment decisions that are less sensitive to vector population abundance. Universal coverage is recommended for ITNs [63], whereas effective vector control intervention by IRS should exceed 30% coverage of households [64] with recommended levels greater than 80% [65]. The ITN covariate shows a geographically broad increasing trend across much of sub-Saharan Africa with values that often exceed 10% in 2020 (Figure S16), whereas non-negligible IRS covariate values are comparatively limited in recent years (Figure S17). Across all samples, the maximum ITN covariate value by grid cell exceeded 10% in 18% of cells with observed genotype data and 86% of cells with observed abundance data. In contrast, the maximum IRS covariate value exceeded 10% in only 3% of grid cells with observed genotype or abundance data. The observational analyses of intervention was therefore restricted to ITNs that exhibited broad geographic deployment of non-negligible interventions. A single abundance study that occurred prior to 2000 was assigned ITN covariate values derived from contemporaneous ITN use data for the study location [66].

### Analysis of Relative Species Abundance

Sinka et al. [27] use multinomial generalised additive models of genotype data to predict relative abundance (or composition) of *An. funestus, An. arabiensis* and the combination of the two species *An. coluzzii* + *An. gambiae s.s*. based on a habitat suitability index [23] and vector intervention data. Here, in contrast to [27], the VectorBase.org data genotyped to species were used to i) model each of the three species *An. arabiensis, An. coluzzii* and *An. gambiae s.s*., which is suggested as a potential avenue for exploration by Sinka et al. [27], and ii) include intra-annually varying predictors for the relative abundances of these species. The analysis included covariates of temperature, precipitation and relative humidity that varied daily, in contrast to previous studies [24, 27–29]. The dates listed in the filtered VectorBase.org genotype data typically included a single day for each record, but some records were assigned a duration of up to a month (see previous section). The meteorological covariates for spatially and temporally varying precipitation, temperature and relative humidity used for the relative abundance analyses were constructed as follows. Thirty day averages were constructed from the daily meteorological observations that preceded both the first and last dates of the collection period for a sample. These two thirty day averages, which coincide for collections with a single day duration, were then averaged together.

ITN use is linked to selective pressure against *An. gambiae s.s*. and in favour of *An. arabiensis* [67, 68] and was also included as a covariate. Collection method was not explicitly included as a covariate. Genotype data for the three species of interest often do not distinguish between outdoor and indoor collection methods [24] that can affect interpretation of intervention efficacy [27]. For example, 27% of genotype samples in this study reported the collection method as “catch of live specimens” (collection method was incorporated into the analysis for the abundance data, see next section). The collected samples represent a mix of indoor and outdoor locations, which may help mitigate biased estimates [27]. Geographical covariates included distances to coasts, elevation, probability of permanent freshwater, human population density censored above 2,000 inhabitant per square kilometre, longitude and the absolute value of latitude as a measure of insolation. The year of collection was included to account for changes in species composition change due to increasing capacity for taxonomic discrimination at the species level for genotype data [69].

Multinomial logit generalised linear models (GLMs), see [70], were analysed using R package **nnet** [71] with the covariates scaled between zero and one based on the observed ranges. Let m ∈ {*Aa, Ac, Ag*} indicate the three species of *An. arabiensis* (*Aa*), *An. coluzzii* (*Ac*) and *An. gambiae s.s*. (*Ag*) with probability *p*_*m*_(*s, t*) at grid cell *s* and time *t*. Let *ν*(*s, t*) denote the set of covariates at grid cell *s* and time *t. An. arabiensis* was chosen as the baseline or reference category of the multinomial logit GLM. The logarithm of the odds *p*_*m*_^*′*^ (*s, t*)*/p*_*Aa*_(*s, t*) in favour of *An. coluzzii* (*m*′ = *Ac*) or *An. gambiae s.s*. (*m*′ = *Ag*) and against *An. arabiensis* were related to the covariates by the unknown parameter vector *γ*. The statistical model is

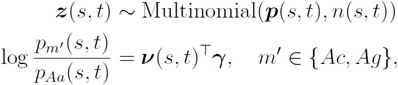

where *z*(*s, t*) is the vector of counts of each species, given by *z*_*m*_(*s, t*) for species *m*, which sums to total observed abundances *n*(*s, t*), and has corresponding probabilities ***p***(*s, t*). The parameter vector *γ* was estimated by maximum likelihood. The following covariates were assessed by model selection. Trends in species composition were assessed by including the annually-varying covariates ITN use and year, which were included as linear terms together with their interaction. The interaction term was included to consider approximate changes in ITN efficacy over time, as could occur during pervasive trends in insecticide resistance that have been observed for *An. gambiae s.l*. in some regions [72]. A quadratic surface model was used as a second order approximation for the remaining meteorological and geographical covariates, that is, each of these covariates appeared as a squared term and a linear term, along with all pairwise interactions of the latter. Backwards stepwise model selection used the R package **MASS** based on the Bayesian information criterion (BIC) [71]. The model with the lowest BIC value was chosen as the final model.

### Analysis of Abundance of *An. gambiae s.l*

The observed abundance *y*(*s, t*) of adult female *An. gambiae s.l*. per person at “site” (grid cell) *s* and time *t* was adjusted to account for different observation methods. The recorded collection methods from the VectorBase.org abundance dataset were mapped to covariates that assigned indoor human landing catch as the baseline method. Binary factors were scored for collection methods of light traps, exit traps, pyrethrum spray catches, animal baited traps and pit traps (Table S4), and whether or not a collection method was documented to occur outdoors. Projects with collection methods that were uncertain with respect to indoor or outdoor sampling were assigned a value of 0.5 for outdoors. The observation method according to Table S4 was encoded by the covariate vector *w*(*s, t, k*) with unknown coefficients *ζ* to adjust the mean abundance per indoor human *f* (*s, t*) by the multiplicative factor *ρ*(*s, t, k*) to produce the observed mean abundance per indoor human *µ*(*s, t, k*) for collection method *k*. If *w*(*s, t, k*) is a zero vector then the corresponding collection method is “man biting catch - indoors” (MBCI, Table S4) with *ρ*(*s, t, k* = MBCI) = 1 and *µ*(*s, t, k* = MBCI) = *f* (*s, t*).

The equilibrium vector population abundance per indoor human *f* (*s, t*) was assumed to be described by the spatially and temporally varying covariates *v*(*s, t*) with unknown coefficients *β*. Human population density is a proxy index for anthropogenic habitat modification and urbanisation. For example, the density of houses is negatively associated with larval habitat area for *Anopheles* mosquitoes [73]. High human population density not only reduces larval habitat but also reduces per capita malaria exposure, suggesting lower vector areal density in urban areas [55, 56]. However, in rural areas, human population is positively associated with *An. gambiae s.l*. areal density [7, 37]. A quadratic relationship on the censored human population density covariate was therefore included in the model. The linear and squared terms allow the potential per capita *An. gambiae s.l*. abundance to increase at low population densities and decrease at high population densities approaching the urban population density threshold. The covariate vector ***v***(*s, t*) also incorporated the lagged effect of precipitation on larval habitat. For example, the carrying capacity for *An. gambiae* is related to weekly total precipitation by [21]. Here, the immature phase of the life cycle was assumed to be about 10 days [37]. Thus, for a given day and location, the precipitation covariate was constructed by first calculating the weekly average precipitation that preceded each day of the corresponding 10-day larval period at that location. The weekly running means were then averaged over these 10 days to define the contribution of precipitation to the carrying capacity experienced by *An. gambiae s.l*. mosquitoes during the immature stage of development. The precipitation covariate was included in the model as a quadratic relationship to allow for increased per capita *An. gambiae s.l*. abundance at low precipitation levels and decreasing per capita *An. gambiae s.l*. abundance with high precipitation. Relative humidity, which varies with temperature and like precipitation is also a function of water vapour in the air, has been suggested to be an important factor for *An. gambiae* abundance [45] and so was additionally included as a covariate; the effect of temperature is further considered in the next section. Linear interaction terms between the censored human population density covariate and the meteorological covariates of precipitation and relative humidity were also included in the model.

The statistical model for vector abundance per indoor human given collection method *k* was described by a negative binomial GLM,

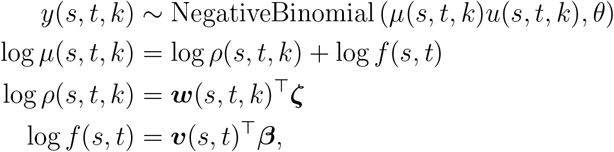

with mean and variance

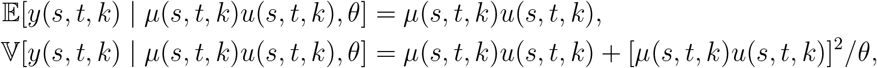

where *θ* is the overdispersion parameter and *u*(*s, t, k*) is an offset for the sampling effort for observation method *k* at cell *s* on day *t* as described above (Methods–Genotype and Abundance Datasets). Unknown parameters were estimated by maximisation of the likelihood with the covariates scaled between zero and one based on their ranges. The negative binomial GLM was fit in R v4.4.0 [74] using the R package **glm2** [75] and function glm.nb in package **MASS** [71].

### Predicted Species Equilibrium Abundance

The equilibrium vector abundances of the three species *An. arabiensis, An. coluzzii* and *An. gambiae s.s*. were predicted for the time period from 1 January 2002 to 31 December 2020 across the potential spatial range of *An. gambiae s.l*. [24, 38]. The predictions of relative abundance for the three species were based on the multinomial logit GLM with the year covariate set to the last year of the prediction period, 2020. The predictions for mean abundance of adult female *An. gambiae s.l*. per human at site (grid cell) *s* and time (day) *t* were based on the negative binomial GLM with the collection method set to indoor human landing catch (MBCI), such that *f* (*s, t*) = *µ*(*s, t, k* = MBCI). Abundance predictions of *An. gambiae s.l*. are best related to precipitation in regions where temperature is not a limiting factor [76]. A temperature threshold rule was therefore applied. Laboratory experiments suggest that larval emergence of *An. gambiae s.l*. does not occur if water temperatures are less than 18°C or more than 34°C [77]. Air temperatures tend to be lower than water temperatures in *Anopheles gambiae s.l*. larval habitats [78], although the exact relationship between water and air temperatures can vary depending on local environmental factors [79]. About 1% of data observations included average air temperatures that were outside the 18^*°*^C – 34°C range. Seasonal reproduction could potentially still occur in such areas during favourable months, but such regions were not represented well enough in the abundance sample data to capture such detail in the predictions. Sites where air temperatures were outside these temperature tolerances for at least 30 continuous days in each year were filtered out to avoid overextrapolation and were provisionally assigned predictions of zero *An. gambiae s.l*. abundance, such that *µ*(*s, t*) = 0 for all *t* at any site *s* that exceeded the temperature tolerance for larval emergence.

The negative binomial GLM provided per capita estimates of adult females for the *An. gambiae* species complex based on counts of *An. gambiae s.l*. Species level predictions therefore require incorporation of the multinomial GLM that provides relative abundance estimates for the three species of interest. A complication is that the predicted mean abundance *f* (*s, t*) = *µ*(*s, t, k* = MBCI) in the negative binomial GLM is the average daily abundance of adult female *An. gambiae s.l*. per human based on indoor feeding mosquitoes, but the propensity for human feeding among the three species of interest is not the same and adult female mosquitoes do not feed every day. Human feeding rates across the species range of *An. gambiae s.l*. have been elicited from domain experts [80] accounting for species differences among *An. arabiensis, An. coluzzii* and *An. gambiae s.s*. using a prior elicitation method for GLMs [81]. Estimates of the average daily probabilities of an adult female mosquito biting a human are ***r*** = [0.37, 0.49, 0.47]^T^ for *An. arabiensis, An. coluzzii* and *An. gambiae s.s*., respectively [80]. These average estimates reflect the greater degree of zoophilic feeding habit of *An. arabiensis* [23], although the human feeding rate will in actuality vary with co-variates such as livestock density. The feeding rate adjustment for total adult females of the *An. gambiae* complex specifies that there are more adult female mosquitoes than those that may be observed feeding on humans, and this effect is greater on average for *An. arabiensis* compared to the other two species. Let *ξ*(*s, t*) denote the average abundance of adult female *An. gambiae s.l*. per human after this adjustment for the mean daily probability of human feeding. Using the predicted per capita abundance *f* (*s, t*) from the negative binomial GLM and the predicted relative abundances ***p***(*s, t*) from the multinomial logit GLM obtains

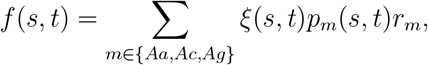

so that *An. gambiae s.l*. vector abundance per human is

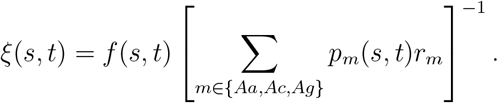

The predicted species vector abundances ***x***(*s, t*) were then obtained by again using the multinomial GLM predictions,

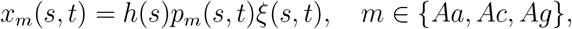

where *h*(*s, t*) is the human population at site *s* and time *t* censored at the urban threshold of 50,000 inhabitants in a 5 km *×* 5 km grid cell. As described above, the total human population is censored because the relationship between vector abundance and human population is weaker in urban environments [55, 56] compared to non-urban areas [7, 37]. Note that the relative abundance of each species based on the total vector abundances is again *p*_*m*_(*s, t*) = *x*_*m*_(*s, t*)*/Σ* _*m*_ *x*_*m*_(*s, t*) as originally predicted by the multinomial GLM.

The multinomial and negative binomial GLMs each accounted for various environmental and human factors that affect the average carrying capacities of the three species. For prediction, the models were interpreted to track potential equilibrium vector abundances determined by the running averages of the meteorological covariates, rather than actual abundances, subject to local conditions not accounted for in the continent-wide spatial and temporal covariates used in this analysis. The predicted equilibrium vector abundance at a given site may not be realised for many important ecological factors not considered in this model, for example, immigration or emigration due to metacommunity dynamics (see [80] for an elaborated model to address these considerations including wind-assisted advection, sensu [82]). The predicted species abundances ***x***(*s, t*) were therefore considered predictions of equilibrium species vector abundances across the various sites and times. An underlying dynamic model that allows for stable coexistence and competing species at equilibrium is presented in the next section.

### Estimated Niche Overlap and Interspecific Competition

Pombi et al. [30] estimate the niche overlap among the three species *An. arabiensis, An. coluzzii* and *An. gambiae s.s*. based on field samples collected at three sites in central Burkina Faso. The pairwise estimates of niche overlap *α*_*ij*_ between species *i* and species *j* use the method of Pianka [31]. For comparison, this method was applied to the equilibrium predictions that are available for all days and grid cells in Burkina Faso for years 2002–2020. Pianka’s niche overlap [31] was estimated from the predicted mean proportion 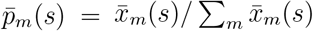 of each species *m* with average vector abundance 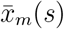 averaged over seasons and years at each site *s* in Burkina Faso to obtain

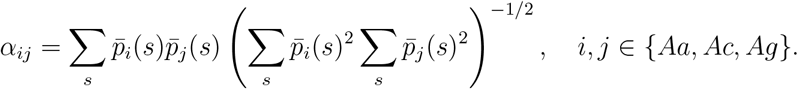

The resulting niche overlap matrix is positive definite and symmetric with unit entries on the main diagonal [32]. The niche overlap index has been suggested as a measure of interspecific competition [32], although Pianka [31] lists limitations with this interpretation.

A contrasting approach estimates interspecific competition by identifying an underlying dynamic model [83] that is a multispecies version of the Beverton-Holt population model [84] or two-species Leslie-Gower model [85] with the added inclusion of a maturation delay. Envision a model for adult female mosquitoes given by

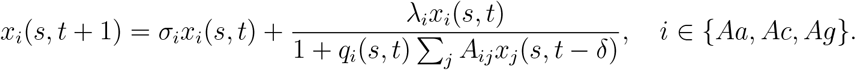

The dynamics of the set of competing species are described at each day *t* with immature life history stages of duration *δ* = 10 days. The dynamic model allows for competitive effects among immature individuals of species *j* on species *i* as mediated by *A*_*ij*_ ≥ 0 that emerge as reproductive adults after the conclusion of the immature life history stage. It is assumed that *A*_*ii*_ = 1 so that strength of intraspecific competition is defined by *q*_*i*_(*s, t*). The density independent per-capita growth rate is *λ*_*i*_. The probability of daily adult survival is *σ*_*i*_. The positive parameter *q*_*i*_(*s, t*) *>* 0 varies through space and time to set the equilibrium level of species *i*.

If ***q***(*s, t*) is held constant, then from [83] the resulting equilibrium is

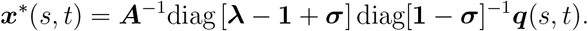

Let ***q***(*s, t*) have spatial and temporal average 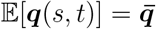. The predicted average equilibrium is then defined by the spatial and temporal average,

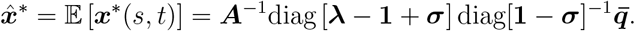

Given the above model, if it is assumed that the average vector abundance 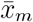 of species *m* over all sites and times is an approximation of its predicted equilibrium vector abundance, such that 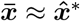, then following [83] a minimisation problem can be constructed that seeks to shift the bottom of the “bowl” formed by the basin of attraction towards the estimated equilibrium 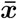 while increasing the magnitude of interspecific competition described by the off-diagonal entries in ***A***. The underlying premise is that species “seek” to maximise available resources, thus increasing competition, and are more likely to be observed near an equilibrium that is stable and more resilient [83]. Stability is measured by the spectral radius, or largest eigenvalue by magnitude, of the linearised system around the equilibrium (see box). If the spectral radius is less than one, then the system is locally stable such that species abundances will eventually return to equilibrium after a small perturbation. As the spectral radius decreases below one, the return time to the equilibrium following a perturbation decreases and the system is said to be more resilient [86]. The symmetric positive definite structure of the niche overlap method is retained for ***A*** in this approach.

#### Methods Box

Understanding Stability and Resilience via the Spectral Radius

The potential for stable coexistence of competing species is assessed by examining how small disturbances behave around the equilibrium of a dynamic system. This mathematical approach uses a linearised approximation of a often nonlinear system to predict how species respond to small changes in abundances near equilibrium. For the nonlinear delayed model considered here, the linear approximation depends on the competition matrix ***A***, equilibrium abundances ***x***^*^, and known fecundity and survival parameters [83]. The diagonal terms in the competition matrix ***A*** determine the strength of intraspecific competition, while off-diagonal terms determine the strength of interspecific competition. The largest eigenvalue of a linearised discrete time system is called the spectral radius that is denoted by *ρ*. The spectral radius determines whether stable coexistence is possible:

*ρ <* 1: the system is locally stable and will return to the equilibrium after a small disturbance, which means the species will persist and coexist.

*ρ >* 1: the system is unstable and may not return to the equilibrium after a small disturbance, which could lead to local species extinctions.

Moreover, the magnitude of the spectral radius determines the resilience of the system that is defined by the asymptotic return time to a stable equilibrium: a smaller spectral radius indicates greater resilience and faster rate of return after a perturbation that increases the chances of persistence and reduces the risk of extinction [86]. In this study, the concepts of stability and resilience defined by the spectral radius are used to evaluate the plausibility of competitive coexistence among *An. arabiensis, An. coluzzii*, and *An. gambiae s.s*.. The strength of competition among species is estimated by selecting parameter combinations that maximise the product of system resilience and interspecific competition given the observed species relative abundances [83]. This approach finds an optimised system where competition for resources and the likelihood to persist under natural environmental variability are both maximised.

The estimate used the ten day immature stage duration (*δ*_*m*_ = 10 days) with fecundity rate of *λ*_*m*_ = 9*/*2 female adults estimated in [37] for *An. gambiae s.l*. and adjusted for equal sex ratio. See also [83] for an example using the empirical results of Pombi et al. [30] for central Burkina Faso only. Adult daily survival probabilities across the species range of *An. gambiae s.l*. were elicited from domain experts [80] accounting for species differences among *An. arabiensis, An. coluzzii* and *An. gambiae s.s*. using a prior elicitation method for GLMs [81]. The average predicted survival probabilities of ***σ*** = [0.85, 0.84, 0.84]^T^ for the species *An. arabiensis, An. coluzzii* and *An. gambiae s.s*., respectively, were obtained from [80]. The estimate of competition does not depend on ***q*** in the method of [83], so that the above assumptions allowed estimation of the interspecific density dependent coefficients *A*_*ij*_ with *i ≠ j* over all sites in Burkina Faso for the period 2002–2020.

## Results

### Relative Abundance and Per Capita Abundance GLMs

The coefficient estimates and standard errors of the best fitting model for the log odds of *An. coluzzii* and *An. gambiae s.s*. against *An. arabiensis* as the baseline or reference category are presented in Table S3. The backwards stepwise selection resulted in a best fitting final model having a BIC value of 49,832 compared to 49,899 for the initial full model. The meteorological covariates of precipitation, temperature and relative humidity were retained as quadratic relationships, with both linear and quadratic terms retained as useful predictors of relative abundance, in the best fitting model. The relative abundances of both *An. coluzzii* and *An. gambiae s.s*. showed a decreasing trend with higher latitudes, though this effect was modified by interactions with temperature, human population density, elevation, distance to coasts and the probability of permanent water. Human population density was evidently an important predictor included as both linear and quadratic terms as well as interactions, where it showed a negative interaction with longitude for the log odds of *An. coluzzii* and *An. gambiae s.s*. The estimates indicated an increasing annual trend in the relative abundance of *An. coluzzii* and *An. gambiae s.s*., but the temporal trends also interacted with ITN use. By the end of the observation period in 2021, the estimated effect of ITN use suggested a suppression of *An. gambiae s.s*. in favour of *An. arabiensis*.

Predictions from the multinomial GLM are averaged through time and presented across the spatial scope in Figure S21. The associated predicted variance on the linear predictor scale is presented in Figure S22. Higher relative abundance was predicted for *An. coluzzii* in areas located along the northwestern and eastern coasts of the Gulf of Guinea, and inland central West Africa and central Africa. The latter region lacked relative abundance data sources (Figure S1) and predictions of higher relative abundance for *An. coluzzii* versus *An. gambiae s.s*. were accompanied by high uncertainty (Figure S22). Greater relative abundance was predicted for *An. gambiae s.s*. versus *An. arabiensis* south of the Sahel in West Africa, eastern coastal Madagascar, and some regions near Lake Victoria. High relative abundance for *An. arabiensis* was predicted in southern Africa, the Sahel, and much of East Africa including the Horn of Africa, Ethiopian highlands and some regions near Lake Victoria.

The parameter estimates for the negative binomial GLM are provided in Table S5. Observed counts were strongly affected by observation method. Alternative methods to indoor human landing catch (MBCI) were generally estimated to produce higher counts per deployment. Both the precipitation and the censored human population density covariates indicated concave relationships with per capita vector abundance (Figure S23). The concave relationship between per capita vector abundance and human population density induced a unimodal relationship between total vector abundance and human population in a 5 km *×* 5 km grid cell. Total adult vector abundance peaked at intermediate human populations and declined towards zero as the human population approached the urban threshold of 50,000 people in a 5 km *×* 5 km grid cell. A positive interaction was estimated between human population density and precipitation, and a negative interaction between human population density and relative humidity (Table S5). ITN use was negatively associated with per capita abundance. The highest overall per capita abundance was predicted near the equator, along the Gulf of Guinea, the Ethiopian Highlands and Madagascar (Figure S24a) with some areas of high uncertainty (Figure S24b) located in regions with comparatively few abundance data sources (Figure S2). These predictions from the negative binomial GLM do not account for temperature tolerances that are addressed in the next section below.

### Predicted Equilibrium Species Abundances

The multinomial and negative binomial GLMs were used to predict the daily equilibrium abundances of adult female *An. arabiensis, An. coluzzii* and *An. gambiae s.s*. in each 5 km *×* 5 km grid cell across the range of *An. gambiae s.l*. in Africa after accounting for temperature tolerances, daily probabilities of human feeding, and the human population. The average equilibrium vector abundances of each species are presented in Figure 1. The predicted equilibrium vector abundance of *An. arabiensis* was highest in eastern Africa and the Sahelian region and Madagascar with low areal densities in central Africa. The predicted equilibrium vector abundance of *An. coluzzii* was highest in western Africa and absent in eastern Africa. The highest predicted equilibrium vector abundance for *An. gambiae s.s*. occurred in western Africa, the Lake Victoria basin and Madagascar.

**Figure 1.**
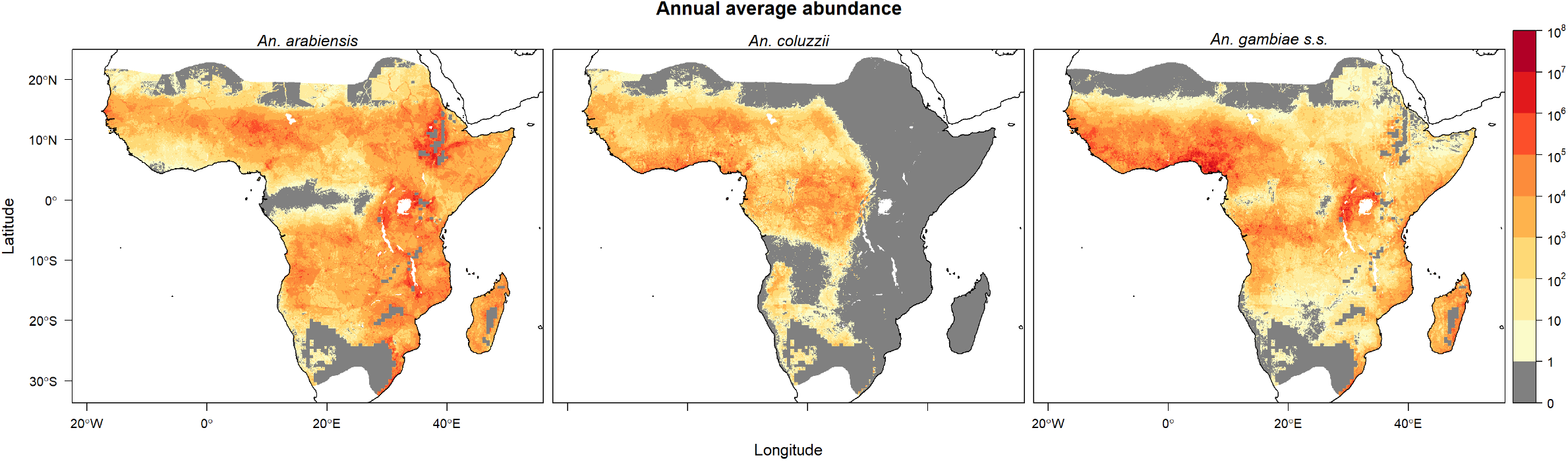
Annual average of predicted equilibrium vector abundances of dominant malaria vector species of *An. gambiae s.l*. per 25 square kilometres over years 2002–2020. Predictions are masked to the potential range of *An. gambiae s.l*. [24] provided by the Malaria Atlas Project (https://malariaatlas.org/) under the terms of a Creative Commons Attribution 4.0 International License available at https://creativecommons.org/licenses/by/4.0/. Note logarithmic scale.

Seasonal predicted equilibrium abundances are visualised in Figures 2, 3 and 4, where seasonally high abundances for all three species occurred during the third quarter (Q3, July– September) in West Africa when precipitation was high (Figure S6). High abundance during this season was also predicted for *An. arabiensis* in regions of the Ethiopian highlands near the limits of the temperature tolerance for *An. gambiae s.l*. The opposite seasonal pattern was predicted in the southern parts of the ranges of *An. arabiensis* and *An. gambiae s.s*., where precipitation is generally higher in the first quarter (Q1, January–March) compared to the third quarter (Figure S6). In these southern regions, high equilibrium abundances were predicted for both *An. arabiensis* and *An. gambiae s.s*. in the first quarter of year.

**Figure 2.**
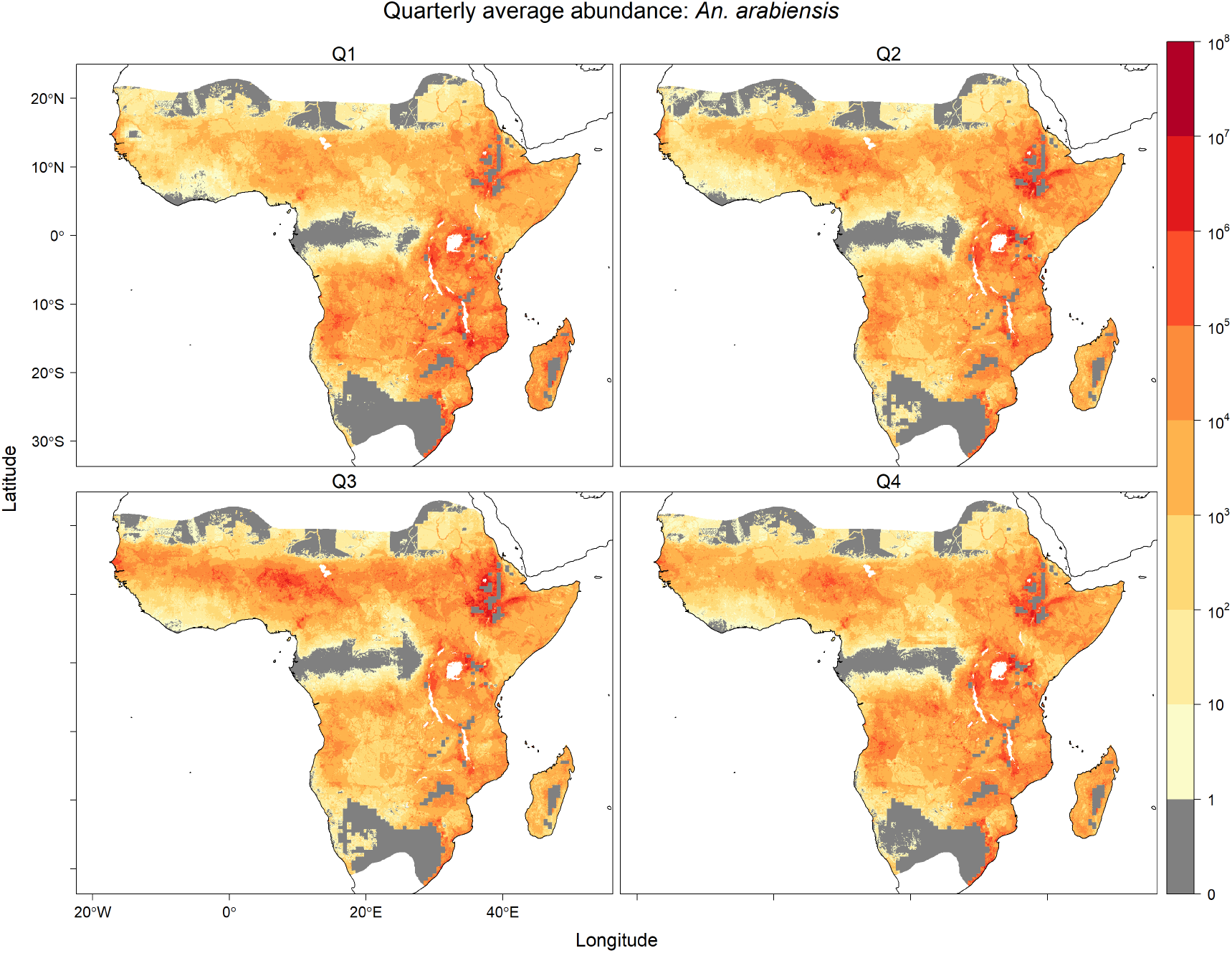
Equilibrium vector abundance of *An. arabiensis* per 25 square kilometres by quarter (Q1: January–March, Q2: April–June, Q3: July–September, Q4: October–December) averaged over years 2002–2020. Predictions are masked to the potential range of *An. gambiae s.l*. [24] provided by the Malaria Atlas Project (https://malariaatlas.org/) under the terms of a Creative Commons Attribution 4.0 International License available at https://creativecommons.org/licenses/by/4.0/. Note logarithmic scale.

**Figure 3.**
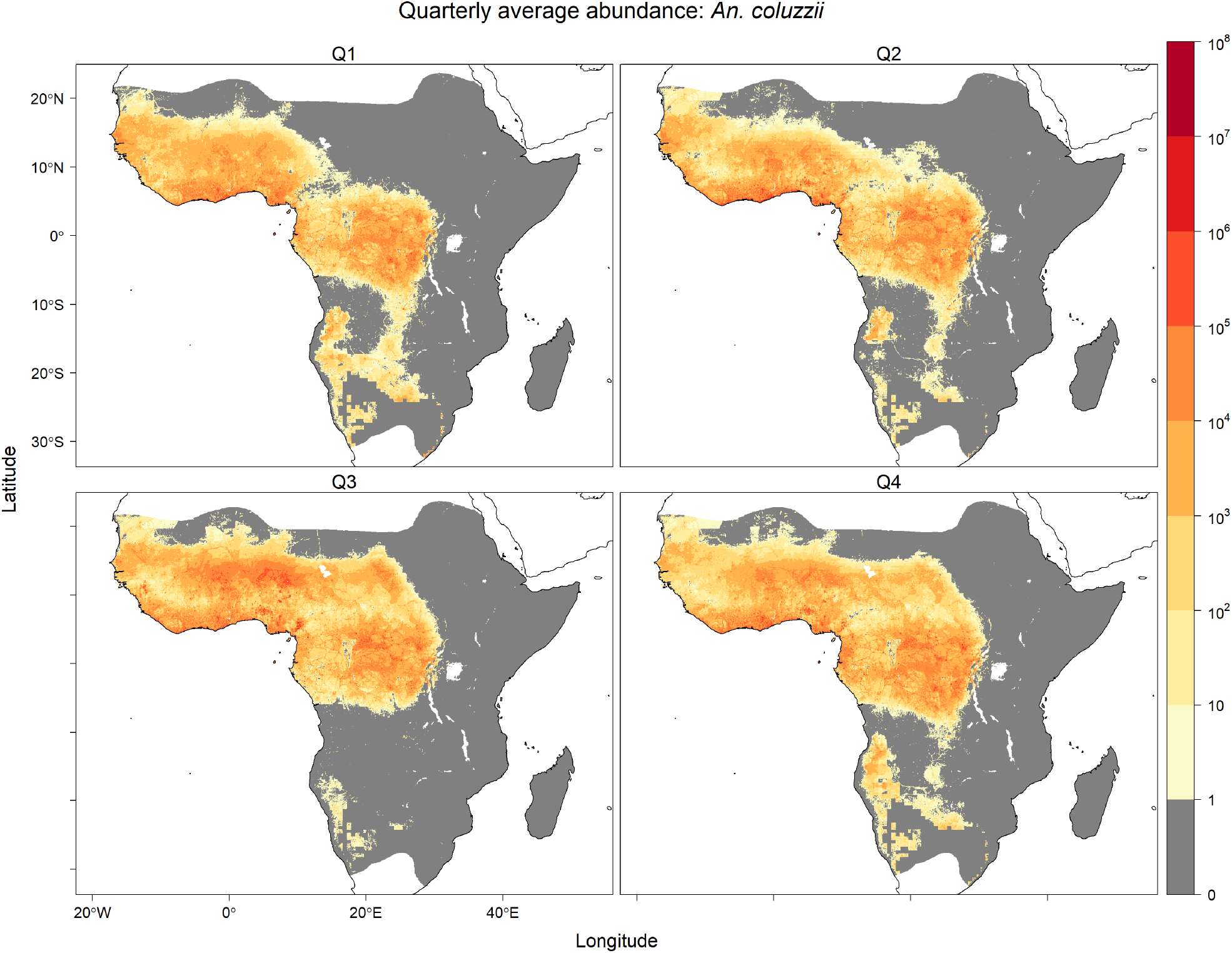
Equilibrium vector abundance of *An. coluzzii* per 25 square kilometres by quarter (Q1: January–March, Q2: April–June, Q3: July–September, Q4: October–December) averaged over years 2002–2020. Predictions are masked to the potential range of *An. gambiae s.l*. [24] provided by the Malaria Atlas Project (https://malariaatlas.org/) under the terms of a Creative Commons Attribution 4.0 International License available at https://creativecommons.org/licenses/by/4.0/. Note logarithmic scale.

**Figure 4.**
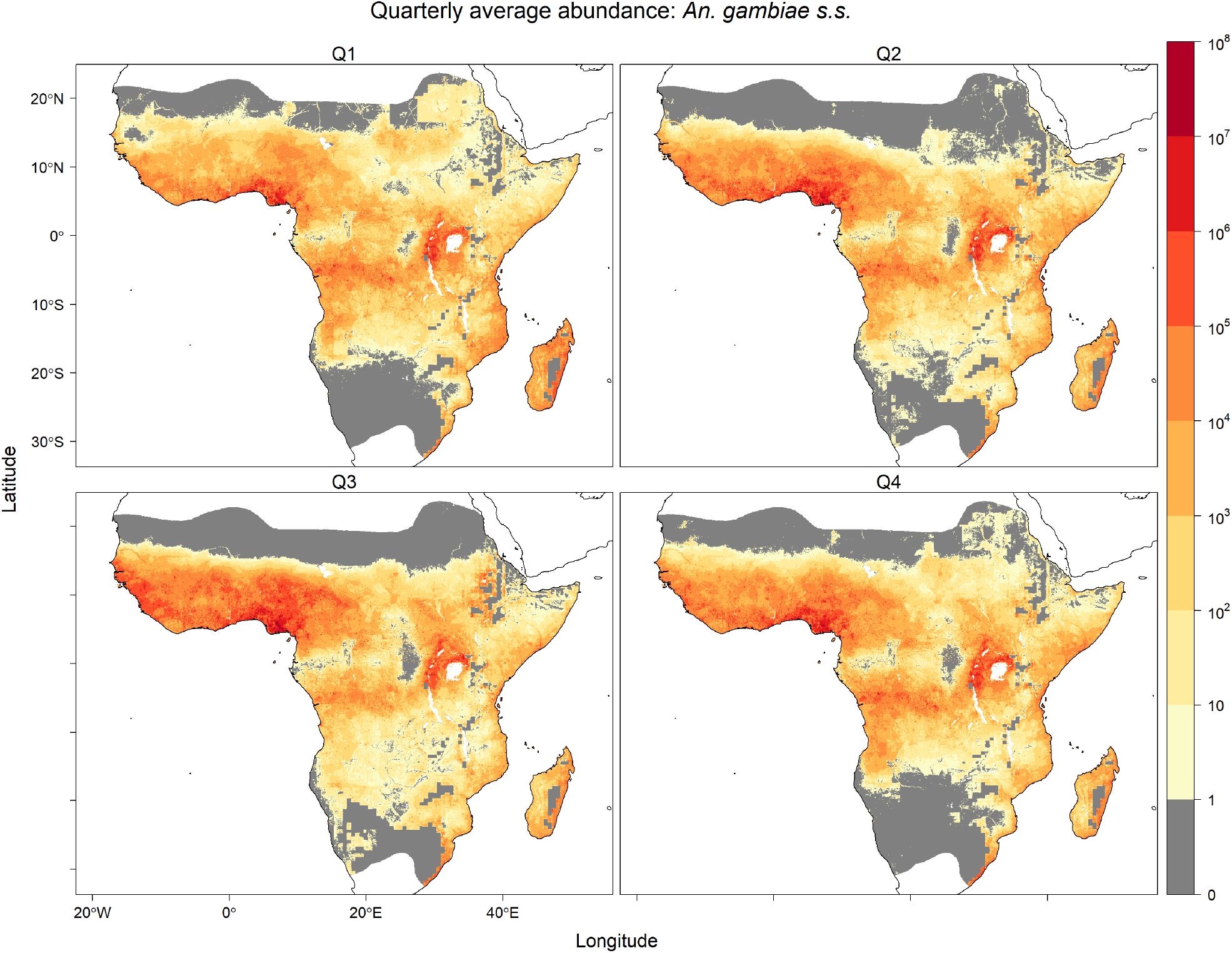
Equilibrium vector abundance of *An. gambiae* s.s. per 25 square kilometres by quarter (Q1: January–March, Q2: April–June, Q3: July–September, Q4: October– December) averaged over years 2002–2020. Predictions are masked to the potential range of *An. gambiae s.l*. [24] provided by the Malaria Atlas Project (https://malariaatlas.org/) under the terms of a Creative Commons Attribution 4.0 International License available at https://creativecommons.org/licenses/by/4.0/. Note logarithmic scale.

Predicted equilibrium abundances for the three species also varied across years as illustrated by continent wide predictions separated by six year intervals between 2002–2020 (Figures S18, S19, S20). The annual average abundance of *An. arabiensis* was generally predicted to be higher in 2002 than in 2020 for most regions outside of the Ethiopian highlands. The annual average abundance of *An. coluzzii* was predicted to be higher in central Africa in 2002 than in 2020, and lower in West Africa in 2002 compared to 2020. Predictions of annual average anomalies for *An. gambiae s.s*. were more similar to *An. arabiensis* than *An. coluzzii*.

### Comparative Evaluation of Equilibrium Abundance Predictions

The predicted daily species equilibrium vector abundances were evaluated by comparison to previous empirical analyses of large spatial scale prediction of species in the *An. gambiae* species complex from four distinct perspectives: i) species distribution models of species occurrence, ii) relative abundance models, iii) observed entomological inoculation rate (EIR) and iv) effective population size estimates from genotype data. Studies in i) and ii) that incorporate species occurrence data may not be entirely independent of the data used for the multinomial GLM model of species relative abundance in this study. However, the comparisons iii) and iv) provide independent empirical comparisons. None of the comparative studies allow for intra-annually varying covariates, and so comparisons considered only the mean equilibrium vector abundance (Figure 1) averaged over daily, and seasonal (e.g., Figures 2, 3 and 4), variation for the 2002–2020 prediction period.

### i) Species Distribution Models

Species distribution models based on species occurrence do not estimate abundance but instead “potential distribution” [26] or the relative probability of presence (e.g., [24]) and so are not directly comparable with this study. Qualitative comparison is still possible for the several species distribution models applied within the *An. gambiae* complex [22, 24, 26, 87, 88]. Of these, only Wiebe et al. [24] predict relative probabilities of presence for all three species based on annually varying covariates. The species distribution model predictions in that study [24] are clipped to independently derived species range limits, but comparison within the limits suggests broad qualitative agreement with the average equilibrium vector abundance predictions of Figure 1: *An. arabiensis* estimates are highest in the northern, eastern and southern regions (see also [22, 26, 88]); *An. coluzzii* estimates are highest in western Africa; and *An. gambiae s.s*. estimates are highest in western Africa and near Lake Victoria with high estimates also occurring in central Africa and the eastern coast of Madagascar. The range limits imposed by Wiebe et al. [24] exclude *An. gambiae s.s*. from the northeastern portion of the range for *An. gambiae s.l*. In this region, Figure 1 shows lower densities extending into the Horn of Africa (the Somali Peninsula and eastern Ethiopia) compared to *An. arabiensis*. Species distribution models that permit predictions into the Horn of Africa suggest either presence [22, 87] or absence [88] of *An. gambiae s.s*.

### ii) Relative Abundance Models

Extrapolations from relative abundance estimates of *An. arabiensis* versus the aggregate of *An. coluzzii* and *An. gambiae s.s*. collated for Nigeria, Niger and the Senegambia in an early study by Lindsay et al. [28] suggest strong similarities with Figure 1. A similar pattern is predicted by Sinka et al. [27]. A discrepancy is noted in southern Tanzania and Mozambique, where the relative abundance of *An. arabiensis* is higher in the two comparative studies than in Figure 1. The extrapolated predictions of the Lindsay et al. study [28] suggest higher relative abundance of *An. gambiae s.s*. in some areas of Ethiopia, unlike the predictions of Figure 1 or the study by Sinka et al. [27] that accord with empirical observations of *An. arabiensis* rather than *An. gambiae s.s*. in this area [89]. Although restricted to comparisons of *An. coluzzii* and *An. gambiae s.s*. relative abundance based on species occurrence data from western and central Africa, Tene Fossog et al. [29] similarly predict higher estimates of the former species across the Sahel and higher estimates of *An. gambiae s.s*. in the West Sudanian savannah. A discrepancy occurs in central Africa, where there is a region of higher abundance predicted for *An. coluzzii* compared to *An. gambiae s.s*. in Figure 1, whereas the reverse is suggested by the predictions of Tene Fossog et al. [29]. As noted previously, the predicted relative abundance in this area is a region of extrapolation in Figure 1 with associated higher uncertainty (Figure S24b) due to the lack of available genotyped samples. Consideration of field collected larvae reared to emergence in the laboratory suggests the presence of both species in central Africa [54].

### iii) Entomological Inoculation Rate

Outside of areas with highly successful vector control and malaria case intervention, Rogers et al. [22] suggest a strong relationship between vector abundance and the entomological inoculation rate (EIR) that is equal to the vector abundance multiplied by the human feeding and sporozoite rates divided by the human population. Proceeding therefore with the predicted species equilibrium vector abundances, where ITN use is incorporated, allows comparison with observations of EIR across East Africa before (years 2000–2010) and after (years 2011–2022) widespread increases in ITN use [90]. For this comparison, EIR is estimated from the predicted species equilibrium vector abundances (Figure 1) averaged over 2002–2020 in each grid cell east of 28^*°*^E longitude and south of 4^*°*^N latitude. Human feeding rates were estimated from the daily feeding probabilities reported in the Methods and the median sporozoite rates *An. gambiae s.l*. reported for both time periods by Msugupakulya et al. [90]. There is substantial variability in these estimates and also the EIR observations [90] that include uncertainty in the sporozoite rate and proportion of EIR attributable to various vector species. Comparisons are therefore based on central 0.80 probability intervals constructed from estimates of the 0.10 and 0.90 quantiles. Using observations of EIR [90], the central 0.80 interval for annualised EIR attributable to *An. gambiae s.l*. in East Africa is (0.15, 175) for the 2000–2010 period and (0, 20) for the 2011–2022 period (see Supplementary Information in [90]). The corresponding 0.80 central intervals for annualised EIR based on the predicted species equilibrium vector abundances were (0, 353) in 2002 and (0, 114) in 2020. It should be noted that occurrences of zero sporozoite rates, which are observed for *An. gambiae s.l*. [90], will underestimate vector abundance based on observed EIR. The equilibrium based estimates of EIR, which assumed median sporozoite rates, were indeed somewhat higher than observed EIR. Despite the requisite assumptions and caveats, the studies suggest overlapping 0.80 central intervals for EIR in each time period with decreasing upper bounds since widespread increases in ITN use.

### iv) Effective Population Size

A pan-African estimate of mean total effective population size for *An. coluzzii* and *An. gambiae s.s*. obtained from allele frequency data is between 1.6 *×* 10^9^ and 3.1 *×* 10^9^ after accounting for seasonal fluctuations that are bounded by a minimum of 3.1 *×* 10^7^ and a maximum of 6.2 *×* 10^9^ [91]. The genetic study uses genotype data collected between 2000 and 2012 with the majority of samples collected since 2009 [92]. In 2011, ITN use was widely increasing in East Africa, as noted in iii) above. Regarding the predicted overall equilibrium vector abundance aggregated over the entirety of the *An. gambiae s.l*. species range across sub-Saharan Africa, multiplying the predicted average daily aggregate vector equilibrium abundances for adult female *An. coluzzii* and *An. gambiae s.s*. by a factor of two to allow for a roughly equal sex ratio suggested a total population average daily estimate that decreased from 8.6 *×* 10^10^ individuals in 2002 to 5.3 *×* 10^10^ individuals in 2014 (Figures S19 and S20 depict the 2002 and 2014 anomalies from Figure 1 for *An. coluzzii* and *An. gambiae s.s*.). Assuming a ratio of total to effective population size of 0.10 [93] suggests an effective population size of 5.3 *×* 10^9^ based on the combined predicted equilibrium abundances of *An. coluzzii* and *An. gambiae s.s*. in 2014, which is outside the estimated range for the mean effective population size although of the same order of magnitude and below the seasonal maximum effective population size estimate.

### Estimated Niche Overlap and Interspecific Competition

In Burkina Faso for the years 2002–2020, *An. gambiae s.s*. was predicted to be the dominant species (Figure 5) compared to the other two species each with approximately 25% species composition. The species with the greatest niche overlap was *An. gambiae s.s*. (Table 1). The estimated competitive interactions in ***Â*** that involved *An. gambiae s.s*. were the strongest and of the same magnitude (Table 1). The competitive effect of *An. gambiae s.s*. on *An. arabiensis* and *An. coluzzii* was 39% greater than the reciprocal effects of the latter species on the former. The equilibrium that maximised resilience was further away from the observed mean abundance 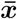 for the Pianka niche overlap estimates (Figure 5a) compared to the optimised competition estimates (Figure 5b). As noted in the Methods, the Pianka index could be considered as a descriptive measure of niche overlap rather than an estimate of competitive interactions [31]. In contrast, for the optimised estimate of interspecific competition (***Â***), the observed mean abundance approximated the most resilient species composition.

**Table 1:** Estimated interspecific competitive interactions in Burkina Faso for the years 2002–2020 using the Pianka niche overlap (*α*) and optimised (***Â***) methods: *An. arabiensis* (Aa) versus *An. coluzzii* (Ac), *An. arabiensis* versus *An. gambiae s.s*. (Ag), and *An. coluzzii* versus *An. gambiae s.s*.

|  | Aa vs Ac | Aa vs Ag | Ac vs Ag |
| --- | --- | --- | --- |
| Pianka | 0.94 | 0.43 | 0.51 |
| Optimised | 0.23 | 0.32 | 0.32 |

**Figure 5.**
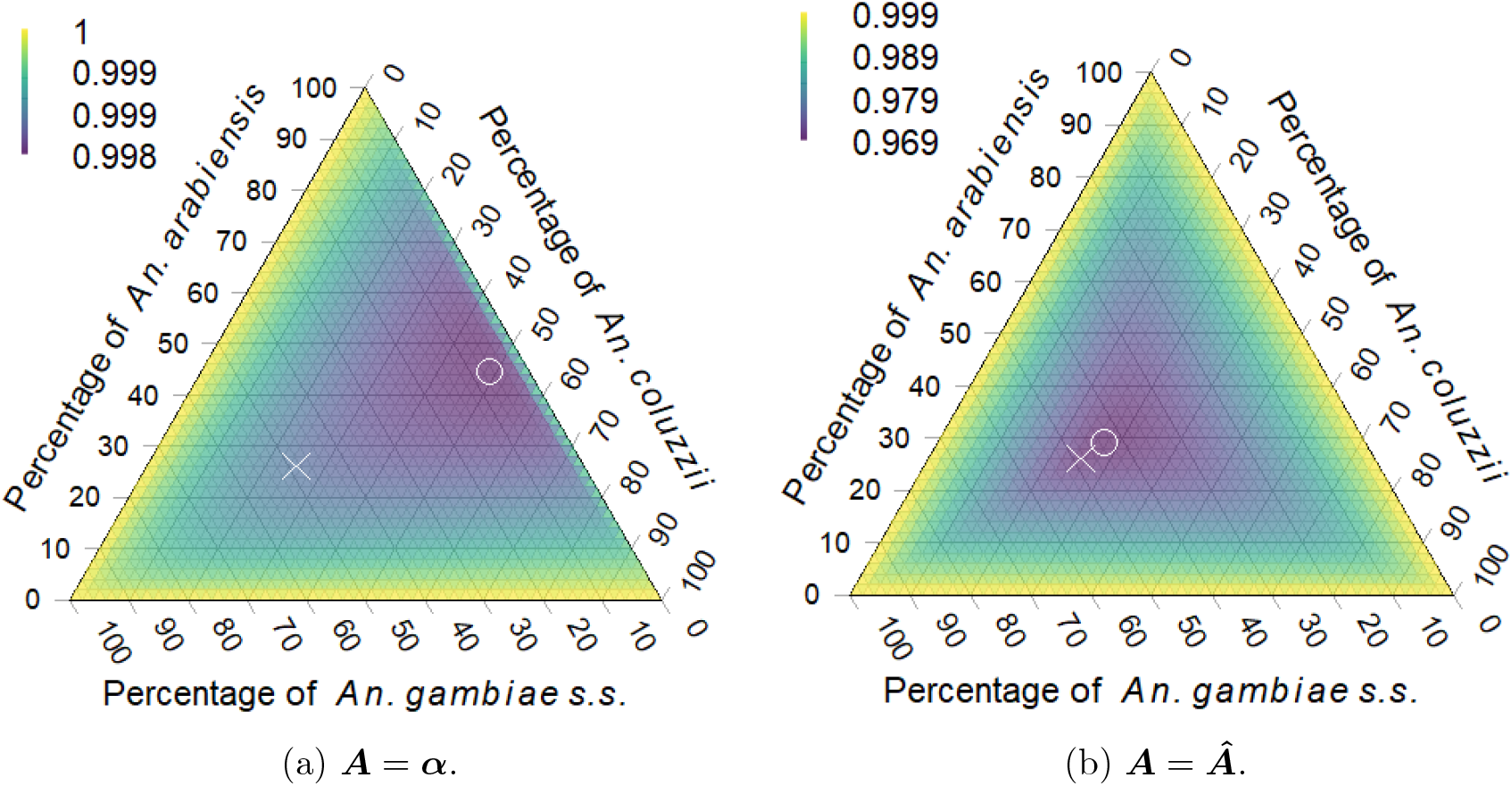
Spectral radius for all possible equilibria where species coexist given (a) Pianka niche overlap estimate ***A*** = ***α*** and (b) optimised estimate ***A*** = ***Â*** for the interspecific competitive interactions. Points show the location of species equilibrium with lowest spectral radius (∘) and the location of the predicted mean equilibrium relative abundances (*×*). Note change in colour scale for (a) versus (b).

## Discussion

Management of malaria requires knowledge of the abundance of important vector species [1, 94] such as members of the *An. gambiae s.l*. species complex [6] to allow efficient malaria control by conventional methods [8] or potential new technologies including genetic biocontrol [7]. The spatiotemporal distribution of vector species abundances, and hence threat of malaria, will change with environmental factors [79] and hence potentially also with climate change [26]. Here, statistically estimated environmental relationships are used to give insights into the predicted abundances of the three important malaria vectors of *An. arabiensis, An. coluzzii* and *An. gambiae s.s*. across their combined ranges of sub-Saharan Africa, and also evaluate the potential for interspecific interactions through niche overlap or competition.

Meteorological factors, such as precipitation and relative humidity, and human factors, such as population density and use of insecticide treated nets (ITNs), were both linked to the relative abundances (species composition) of *An. arabiensis, An. coluzzii* and *An. Gambiae s.s*. and the per capita abundance of *An. gambiae s.l*. These analyses of species relative abundance from genotyped species occurrence data and abundance of *An. gambiae s.l*. from count data were combined to produce joint predictions of daily vector equilibrium abundances for the years 2002–2020 (Figures 1). Predictions showed marked seasonal changes in West Africa for predicted equilibrium abundances of all three species (Figures 2, 3, 4), and a latitudinal progression for *An. arabiensis* and *An. gambiae s.s*. abundances that tracked the wet seasons of higher latitudes. The seasonal variation is explained by the intra-annually varying meteorological covariates of temperature, relative humidity and precipitation.

Predicted equilibrium abundances also varied among years (Figures S18, S19, S20), where for example abundances of *An. arabiensis* and *An. gambiae s.s*. were higher across much of the species ranges in 2002 compared to 2020. Many factors contributed to interannual variation. During this time period, interannual variability in meteorological conditions occurred with more cold days and precipitation for some regions in East Africa, for example, at the end of the 2002–2020 prediction period compared to the beginning [95, 96]. Human population density also increased overall with increased urbanisation and built-up areas [97], while ITN coverage by household in sub-Saharan Africa increased from 5% in 2000 to 70% in 2022 [3]. The equilibrium abundance of *An. gambiae s.l*. was predicted to peak at intermediate levels of human population (Figure S23), and ITN coverage was linked with decreased *An. gambiae s.l*. per capita abundance (Table S5). The relative abundances of *An. coluzzii* and *An. gambiae s.s*. versus *An. arabiensis* were linked to a negative interaction between ITN coverage and year of sample (Table S3). It is possible that the negative interaction between year and ITN coverage for *An. coluzzii* and *An. gambiae s.s*. may be linked to insecticide resistance that is developing with time for some insecticides in certain regions [72], but interpretations of ITN efficacy must be balanced against the observational nature of this study. For *An. coluzzii*, the interaction was indeed small relative to positive estimates for temporal trend and ITN coverage, such that *An. coluzzii* would be predicted to increase across years at the expense of *An. arabiensis* if all other covariates were held constant. The trend of increasing *An. coluzzii* relative abundance could be caused in part to climatic or land use factors, but it could also be explained by the relatively recent species characterisation of *An. coluzzii* and *An. gambiae s.s*. combined with improved genotyping capabilities of the *An. gambiae* species complex [29, 30, 42]. For example, the occurrence of *An. coluzzii* in northwestern Kenya has recently been shown using whole genome sequencing of historical samples dated from 2006 and 2019, where taxonomic characterisation based on previous molecular methods limited to identification of alternative species would have missed detection of *An. coluzzii* [69]. The multinomial GLM estimates (Table S3) suggested that the relative abundance of *An. gambiae s.s*. would generally decrease compared to *An. arabiensis* in later years with high ITN coverage. Temporal shifts in species composition away from *An. gambiae s.s*. towards *An. arabiensis* in areas with high ITN use has been attributed to the outdoor feeding and resting behaviour of the latter species [6, 67, 98].

The predicted equilibrium vector abundance at a given site is a potential abundance that may not be realised for many important ecological factors not considered in this model due to local conditions not accounted for in this continental scale analysis. Predictions based on the model assumptions were therefore comparatively evaluated against four alternative empirical modelling perspectives with general agreement: i) species distribution models based on species occurrence records [22, 24, 26, 87, 88], ii) relative abundance models [27–29], iii) entomological inoculation rate (EIR) estimates in eastern and southern Africa [90] and iv) effective population size estimates from genotype data [91]. Overall, results showed qualitatively similar patterns, although it should be noted that none of the comparisons in i)-ii) provide species-specific estimates of vector abundances. The comparisons iii) and iv), however, provide independent empirical estimates of EIR and vector abundances that are not dissimilar from estimates based on predicted species vector equilibrium abundances.

Analyses suggested that the three species can coexist (Figure 5) despite evidence of substantial niche overlap and interspecific competition (Table 1) across the shared species range. The estimates for niche overlap among the three species (Table 1) for Burkina Faso during 2002–2020 followed a similar pattern to those reported by Pombi et al. [30] in a localised study that derived the estimates from field observations at three sites in central Burkina Faso where all three species coexist. The niche overlap estimates from this study also included the southern region of the country where *An. gambiae s.s*. predominates [99]. Both studies estimated the highest Pianka niche overlap indices between *An. arabiensis* and *An. coluzzii* and similar niche overlaps for comparisons that included *An. gambiae s.s*. (Table 1). The estimates of interspecific competition that maximised interspecific competition and resilience suggested substantial competition was possible in Burkina Faso (Table 1), where all three species occur.

Statistical studies that develop dynamic population models for anophelines from time series data in Africa have targeted species complexes such as *An. gambiae s.l*. or *An. funestus* [20, 21, 100]. Such studies would additionally benefit from geographically and temporally extensive time series data at the species level. Ideally, daily times series of species-level abundances collected over long durations would be available from many sites geographically dispersed across sub-Saharan Africa. Then delayed density dependent models, such as described in the Methods, could be implemented within a state space modelling framework to estimate the parameters of interest while allowing for non-equilibrium dynamics. Without such time series data, this study used smoothed moving averages of the meteorological covariates to approximate daily variation among environmental carrying capacities and equilibrium abundances of the species based on maximum likelihood estimates. The predicted equilibrium is spatially and temporally varying due to changing environmental conditions that modify the Grinellian niche of the constituent species [101], which in turn is modified by interspecific competition to produce the realised niche [102]. However, the predicted equilibrium is only a measure of potential abundance: the species may be limited by dispersal capability or source-sink metapopulation dynamics such that it is either above or below the predicted equilibrium abundance over the long run [11, 103–107]. Moreover, species abundances do not exactly attain potential equilibrium due to changing environment, dispersal and immigration. The proposed density dependent model may be elaborated to account for immigration and emigration, see [12] for a non-delayed example, as well as delayed maturation with introgression among the constituent species of a species complex such as *An. gambiae s.l*. [80].

Given the sparse data available for this study across the broad spatial and temporal scope of interest, and predicted changes of species abundance both within and among years, sophisticated analyses for causal attribution among cumulative effects were commensurately limited. Higher quality longitudinal abundance data at the species level of sufficient sub-seasonal temporal frequency would greatly help statistical evaluation of competing causal hypotheses, such as vector control, malaria interventions, land use and climate. The predictions and results of this study accounted for intra-annual shifts in species composition, but could not explicitly control for many important covariates. For example, factors such as land use impact not only larval habitat but also the feeding habit of *An. arabiensis* that feeds on livestock to a greater degree than *An. coluzzii* and *An. gambiae s.s*. [23]. Inclusion of spatiotemporal covariates of livestock density in particular would assist predictions of *An. arabiensis* that exhibits greater zoophily biting habit. Improved estimation of the relationship between species abundances and human feeding rates would also require abundance data for each of the three species, whereas most of the spatially extensive and longitudinal data available for this study were limited to counts for the entire species complex. Although the ecologically important and spatially and temporally varying covariate of livestock density was not explicitly represented, the lower probability of human host feeding by *An. arabiensis* compared to the other species was incorporated into the predictions. The abundance estimates also accounted for collection methods, but the analysis of the relative abundance data did not incorporate sampling effects because species identification by genotyping were often based on samples collected from the field by multiple methods that did not document, for example, indoor versus outdoor collection. Hence, predictions of *An. arabiensis* relative abundance must be interpreted cautiously and particularly so in regional areas with high livestock density relative to human inhabitant density.

The modelling approach used here elaborated previous relative abundance models [27–29] to consider the three species of *An. arabiensis, An. coluzzii* and *An. gambiae*. The daily predictions of equilibrium abundance at the continental scale for the years 2002–2020 are only approximations of potential mosquito abundance at any given place and time. Daily varying meteorological covariates were used to predict the relative abundances of these species and also the per capita vector abundance of *An. gambiae s.l*. across sub-Saharan Africa for the first time. These statistical models used covariate and response data collated to a grid of 5 km *×* 5 km cells for analysis and prediction at the continental scale. Other regional and continental scale studies use this same resolution [23, 24, 27, 29, 54] as a suitable choice for the ecology and dispersal of species in the *An. gambiae* species complex [23, 29, 54]. At finer resolutions, meteorological factors such as precipitation and temperature can for some locales be more important predictors of *An. gambiae s.s*. abundance counts than microscale covariates such as the number of occupants per home or distance to larval habitat [108]. Climatological factors remain important predictors for resolutions that are coarser than the choice of 5 km *×* 5 km cells [26, 28]. The choice of 5 km *×* 5 km resolution, however, does not deny the known importance of microscale habitat variability to the risk of malaria transmission [73], and is more suitable for interpretation at coarser national or sub-national levels [24].

## Conclusion

Meteorological factors, such as precipitation and relative humidity, and human factors, such as population density and insecticide treated net coverage, were found at the continental scale to be important spatially and temporally varying predictors of vector equilibrium abundances of the primary malaria vector species within the *Anopheles gambiae s.l*. species complex: *An. arabiensis, An. coluzzii* and *An. gambiae s.s*. Daily varying meteorological covariates such as precipitation and relative humidity led to substantial variation in the predicted seasonal equilibrium abundances of the three species. ITN use was negatively associated with per capita vector abundance of the *An. gambiae* species complex. An interaction with ITN use and year predicted a net positive estimate on the ratio of *An. gambiae s.s*. versus *An. arabiensis* around 2002, but this prediction shifted to a net negative against *An. gambiae s.s*. around 2020. A longer term trend of increasing relative abundance of *An. coluzzii* was also suggested, and was possibly attributable to advances in genetic methods of species identification within the *An. gambiae* species complex that occurred over the observational period. Human population density was also an important predictor that suggested a negative associated with per capita vector abundance approaching the threshold for urban areas. Across sub-Saharan Africa, the observational analyses suggested evidence for niche overlap and competitive interactions among the three species. Additional time series data of mosquito vector species abundance, collected at fine temporal resolutions of monthly sampling intervals or higher frequencies and across broad geographical regions, are necessary to improve discernment of ecological interactions among species and the effectiveness of vector control interventions designed to measurably reduce the burden of mosquito-borne disease.

## Supplementary information

The reproducible code, data sources and licenses are summarised in the R package **AgslPredict** [39] provided as a supplement for this article.

## Acknowledgements

The authors thank Scott Foster, Skipton Woolley and the anonymous reviewers for helpful comments. This article incorporates elicited information provided by the following scientific experts: Adedapo Adeogun, Nigerian Institute of Medical Research, Yaba, Lagos, Nigeria; Guillaume Carissimo, Agency for Science, Technology and Research (A^*^STAR), Singapore; George Christophides, Imperial College London, London, UK; Carlo Costantini, Institut de Recherche pour le Développement (IRD), Montpellier, France; Ibrahima Dia, Institut Pasteur de Dakar, Dakar, Senegal; Mawlouth Diallo, Institut Pasteur de Dakar, Dakar, Senegal; Wamdaogo Moussa Guelbeogo, Centre National de Recherche et de Formation sur le Paludisme, Ouagadougou, Burkina Faso; Stephen Higgs, Kansas State University, Manhattan, Kansas, USA; Charles M. Mbogo, KEMRI Centre for Geographic Medicine Research–Coast, Kenya; Juma Mcha, Malaria Elimination Program, Zanzibar, United Republic of Tanzania; Givemore Munhenga, National Institute for Communicable Diseases and University of Witwatersrand, Johannesburg, South Africa; Martin Rono, KEMRI Centre for Geographic Medicine Research–Coast, Kilifi, Kenya; N’fale Sagnon, Centre National de Recherche et de Formation sur le Paludisme, Ouagadougou, Burkina Faso; Dana L. Vanlandingham, Kansas State University, Manhattan, Kansas, USA. The authors thank the experts for the generous contribution of their domain expertise.

## Declarations

### Ethics approval and consent to participate

This article incorporates elicited information with participants providing their consent to participate under ethics approval by the CSIRO Social Science Human Research Ethic Committee (approval number 136/19).

### Consent for publication

The authors provide their consent for publication.

### Data availability

The data sources and licenses are summarised in the R package **AgslPredict** [39] provided as a supplement for this article.

### Materials availability

Not applicable.

### Code availability

The reproducible code is provided in the R package **AgslPredict** [39] provided as a supplement for this article.

### Author contribution

**G. R. Hosack:** Conceptualization, data curation, formal analysis, funding acquisition, investigation, methodology, project administration, software, visualization, writing–original draft, writing–review and editing. **M. El-Hachem:** Conceptualization, data curation, investigation, methodology, software, validation, visualization, writing–review and editing. **A. Ickowicz:** Conceptualization, investigation, methodology, writing–review and editing. **N. J. Beeton:** Conceptualization, investigation, visualization, writing–review and editing. **A. Wilkins:** Conceptualization, writing–review and editing. **K. R. Hayes:** Conceptualization, funding acquisition, visualization, writing–review and editing. **S. S. C. Rund:** Data curation, writing–review and editing. **S. A. Kelly:** Data curation, writing–review and editing. **M. A. McDowell:** Data curation, writing–review and editing.

