## Supplementary Information for "Prediction of mosquito vector abundance for three species in the *Anopheles gambiae* complex"

### Supplementary Information for "Predicted abundance of dominant mosquito-borne disease vectors in *Anopheles gambiae* sensu lato"

Geoffrey R. Hosack[1], Maud El-Hachem[1], Adrien Ickowicz[1], Nicholas J. Beeton[1], Andrew Wilkins[2], Keith R. Hayes[1], Samuel S. C. Rund[3], Sarah A. Kelly[4], and Mary Ann McDowell[3]

[1]CSIRO Data61, 3 Castray Esplanade, Hobart, 7001, Tasmania, Australia
[2]CSIRO Mineral Resources, 1 Technology Court, Pullenvale, 4069, Queensland, Australia
[3]Galvin Life Science Center, University of Notre Dame, Notre Dame, 46556, Indiana, USA
[4]Department of Life Sciences, Imperial College London, London, SW7 2AZ, UK

### Supplementary Tables

Table S1: Data source table for projects from VectorBase.org included in the analysis of species relative abundances.

| Project | Citation |
|---|---|
| VBP0000005 | |
| VBP0000028 | DOI:10.1186/1756-3305-5-69 |
| VBP0000614 | DOI:10.1186/1475-2875-12-24 |
| VBP0000613 | DOI:10.1371/journal.pone.0081931 |
| VBP0000774 | |
| VBP0000009 | |
| VBP0000224 | DOI:10.1186/s12936-017-1756-5 |
| VBP0000624 | DOI:10.1186/1475-2875-13-27 |
| VBP0000148 | DOI:10.1186/s12936-015-0993-8. |
| VBP0000298 | DOI:10.1186/s13071-018-2824-6 |
| VBP0000345 | DOI:10.1101/530758 |
| VBP0000615 | DOI:10.1016/j.actatropica.2010.08.017 |
| VBP0000652 | DOI:10.1186/1475-2875-8-123 |
| VBP0000006 | DOI:10.1186/1472-6785-9-16 |
| VBP0000712 | DOI:10.1093/trstmh/traa113 |
| VBP0000619 | DOI:10.1186/1475-2875-10-151 |
| VBP0000163 | DOI:10.1038/nature24995 |
| VBP0000008 | DOI:10.1186/1472-6785-9-17 |
| VBP0000188 | DOI:10.1093/jme/tjw228 |
| VBP0000785 | |
| VBP0000321 | DOI:10.1371/journal.pone.0203529 |
| VBP0000324 | DOI:10.1186/s13071-018-2857-x |
| VBP0000782 | |
| VBP0000795 | DOI:10.4269/ajtmh.20-1357 |
| VBP0000802 | DOI:10.3389/fpubh.2021.649672 |
| VBP0000141 | DOI:10.1186/s13071-016-1420-x |
| VBP0000786 | |
| VBP0000684 | DOI:10.1186/1475-2875-12-350 |
| VBP0000717 | DOI:10.1186/s13071-019-3523-7 |
| VBP0000153 | DOI:10.1186/s13071-016-1758-0 |
| VBP0000270 | DOI:10.1186/s13071-018-2702-2 |
| VBP0000191 | DOI:10.12688/wellcomeopenres.10662.1 |
| VBP0000186 | PMID:17253060 |
| VBP0000683 | DOI:10.12688/wellcomeopenres.16064.1 |
| VBP0000812 | DOI:10.1073/pnas.2104282119 |
| VBP0000140 | DOI:10.1186/s12879-016-1542-y |
| VBP0000616 | DOI:10.1186/1471-2334-12-275 |
| VBP0000763 | DOI:10.1051/parasite/2021003 |
| VBP0000203 | DOI:10.1186/s12936-017-2099-y |
| VBP0000276 | DOI:10.1186/s12936-018-2293-6 |
| VBP0000780 | https://www.niaid.nih.gov/research/east-africa-international-center-excellence-malaria-research |
| VBP0000004 | DOI:10.1093/molbev/msr199 |
| VBP0000781 | https://www.niaid.nih.gov/research/east-africa-international-center-excellence-malaria-research |
| VBP0000267 | DOI:10.1371/journal.pmed.1002167 |
| VBP0000010 | |
| VBP0000622 | DOI:10.1186/1475-2875-7-231 |

Table S2: Data source table for projects from VectorBase.org included in the analysis of abundance.

| Projects | Citations |
|---|---|
| VBP0000614 | DOI:10.1186/1475-2875-12-24,DOI:10.1186/1475-2875-9-187 |
| VBP0000774 | |
| VBP0000624 | DOI:10.1186/1475-2875-13-27 |
| VBP0000686 | DOI:10.1111/j.1948-7134.2011.00179.x,DOI:10.1111/mve.12084 |
| VBP0000620 | DOI:10.1186/1475-2875-7-138 |
| VBP0000714 | DOI:10.1186/s13071-020-04403-9 |
| VBP0000782 | |
| VBP0000786 | |
| VBP0000621 | DOI:10.1111/j.1365-2915.2006.652.x |
| VBP0000626 | DOI:10.1186/1471-2334-10-119 |
| VBP0000616 | DOI:10.1186/1471-2334-12-275,DOI:10.1186/1756-3305-6-10 |
| VBP0000812 | DOI:10.1073/pnas.2104282119 |
| VBP0000770 | |
| VBP0000701 | DOI:10.1186/1475-2875-13-111 |
| VBP0000627 | DOI:10.1371/journal.pmed.1002167 |
| VBP0000649 | DOI:10.1186/s12936-019-3076-4 |
| VBP0000780 | https://www.niaid.nih.gov/research/east-africa-international-center-excellence-malaria-research |
| VBP0000267 | DOI:10.1371/journal.pmed.1002167 |

Table S3: Multinomial logit GLM for relative abundance based on covariates that are normalised between zero and one based on the minimum and maximum observed values at grid cell centroids: `lon.centroid`, longitude; `lat.centroid`, absolute value of latitude; `ppw`, probability of permanent freshwater; `elev`, elevation; and running means of precipitation (`PRECTOTCORR1m`), temperature (`T2M1m`) and relative humidity (`RH2M1m`). The following covariates were included as linear terms: `origin.year`, year corresponding to start date of sample collection; `ITN`, percent use of insecticide treated nets; and `hpd`, human population density censored above 2,000 inhabitants per square kilometre. See main text for further information on description and definitions of covariates.

| | coeffAcVsAa | coeffAgVsAa | seAcVsAa | seAgVsAa |
|---|---|---|---|---|
| (Intercept) | 72.31 | 30.59 | 7.73 | 3.14 |
| lon.centroid | -12.74 | -18.11 | 12.75 | 3.58 |
| lat.centroid | -187.51 | -45.42 | 12.68 | 3.61 |
| ppw | -8.67 | -19.29 | 5.59 | 2.44 |
| elev | -2.25 | 0.81 | 13.56 | 4.21 |
| d2c | -36.01 | 4.45 | 5.60 | 2.36 |
| PRECTOTCORR1m | 80.56 | 127.86 | 9.16 | 8.74 |
| T2M1m | 24.14 | -7.23 | 14.22 | 7.37 |
| RH2M1m | -81.78 | -41.52 | 6.10 | 3.05 |
| hpd | 39.35 | 60.57 | 4.96 | 3.26 |
| I(lon.centroid^2) | -164.15 | 9.40 | 9.23 | 1.33 |
| I(lat.centroid^2) | 85.22 | 6.97 | 11.94 | 2.05 |
| I(ppw^2) | -1.28 | -1.74 | 0.69 | 0.26 |
| I(elev^2) | -66.44 | -5.72 | 7.11 | 2.05 |
| I(hpd^2) | -10.32 | -10.82 | 0.73 | 0.46 |
| I(PRECTOTCORR1m^2) | 2.05 | 28.13 | 6.93 | 5.59 |
| I(T2M1m^2) | -63.23 | 9.35 | 11.66 | 4.28 |
| I(RH2M1m^2) | 9.01 | 21.02 | 3.38 | 2.04 |
| origin.year | 0.66 | 1.22 | 0.88 | 0.26 |
| ITN | 4.58 | 2.86 | 1.32 | 0.32 |
| lon.centroid:lat.centroid | 53.55 | 47.01 | 15.53 | 2.26 |
| lon.centroid:elev | 63.77 | -22.57 | 16.48 | 2.26 |
| lon.centroid:d2c | 45.80 | -10.47 | 7.57 | 1.33 |
| lon.centroid:PRECTOTCORR1m | -63.34 | -14.35 | 9.34 | 2.32 |
| lon.centroid:T2M1m | 15.07 | -27.36 | 13.33 | 2.94 |
| lon.centroid:RH2M1m | 54.01 | 4.56 | 8.08 | 1.98 |
| lon.centroid:hpd | -14.67 | -16.11 | 4.95 | 1.58 |
| lat.centroid:ppw | -7.87 | 9.45 | 2.92 | 1.10 |
| lat.centroid:elev | 112.80 | 8.12 | 13.50 | 3.82 |
| lat.centroid:d2c | 3.42 | -16.10 | 6.98 | 2.76 |
| lat.centroid:T2M1m | 102.14 | -15.95 | 17.80 | 4.49 |
| lat.centroid:hpd | -4.47 | -32.33 | 5.24 | 2.31 |
| ppw:elev | 11.94 | 13.72 | 5.20 | 1.23 |
| ppw:d2c | 2.20 | -7.94 | 1.54 | 0.64 |
| ppw:PRECTOTCORR1m | -10.41 | -9.61 | 2.63 | 1.44 |
| ppw:T2M1m | -0.66 | 23.30 | 5.16 | 2.41 |
| ppw:RH2M1m | 15.92 | 7.39 | 3.22 | 1.50 |
| ppw:hpd | 9.62 | 8.58 | 0.76 | 0.54 |
| elev:d2c | 9.48 | 18.03 | 4.92 | 1.24 |
| elev:PRECTOTCORR1m | 87.33 | -14.95 | 10.23 | 3.54 |
| elev:T2M1m | -123.17 | -5.42 | 16.21 | 5.02 |
| elev:hpd | 3.23 | -27.60 | 3.17 | 1.72 |
| d2c:PRECTOTCORR1m | -75.05 | -21.14 | 6.45 | 3.17 |
| d2c:RH2M1m | 50.88 | 10.47 | 4.50 | 2.17 |
| d2c:hpd | -14.91 | 1.27 | 1.87 | 0.96 |
| PRECTOTCORR1m:T2M1m | 9.19 | -79.49 | 9.54 | 6.82 |
| PRECTOTCORR1m:RH2M1m | -52.60 | -90.02 | 13.07 | 8.58 |
| T2M1m:RH2M1m | 43.94 | 26.80 | 6.47 | 3.03 |
| T2M1m:hpd | -42.44 | -54.29 | 4.41 | 2.76 |
| RH2M1m:hpd | -0.52 | -7.77 | 2.17 | 1.24 |
| origin.year:ITN | -0.39 | -4.54 | 1.95 | 0.49 |

Table S4: **Coding for collection methods reported in VectorBase.org abundance data.** Collection methods were scored 1 if occurring outdoors and 0 if occurring indoors. Collection methods described by VectorBase.org are grouped by row: *ABC*, animal baited catch ("animal baited net trap catch" or "animal biting catch - outdoors"); *APS*, "artificial pit shelter"; *LTC*, light trap catch ("CDC light trap catch" or "CDC miniature light trap model 512"); *ETC*, "exit trap catch"; *ILTC*, "indoor light trap catch"; *MBC*, "man biting catch"; *MBCI*, "man biting catch - indoors"; *MBCO*, "man biting catch - outdoors"; *OLTC*, "outdoor light trap catch"; and *PSC*, "pyrethrum spray catch". The collection method are scored by column with respect to light traps (*light*), exit traps (*exit*), pyrethrum spray catch (*PSC*), animal baited catch (*animal*), pit shelter (*pit*) and outdoor deployment (*outdoor*). The latter category is scored at an intermediate value of 0.5 where the collection method was indeterminate with respect to indoor versus outdoor sampling. One project that had "CDC light trap" as the listed collection method was scored indoors using the methods description from the original study (67).

| Method | Light | Exit | PSC | Animal | Pit | Outdoor |
|---|---|---|---|---|---|---|
| ABC | 0 | 0 | 0 | 1 | 0 | 1.0 |
| APS | 0 | 0 | 0 | 0 | 1 | 1.0 |
| ETC | 0 | 1 | 0 | 0 | 0 | 0.0 |
| ILTC | 1 | 0 | 0 | 0 | 0 | 0.0 |
| LTC | 1 | 0 | 0 | 0 | 0 | 0.5 |
| MBC | 0 | 0 | 0 | 0 | 0 | 0.5 |
| MBCI | 0 | 0 | 0 | 0 | 0 | 0.0 |
| MBCO | 0 | 0 | 0 | 0 | 0 | 1.0 |
| OLTC | 1 | 0 | 0 | 0 | 0 | 1.0 |
| PSC | 0 | 0 | 1 | 0 | 0 | 0.0 |

Table S5: Negative binomial GLM given estimated overdispersion parameter of 0.28. Collection methods are described in Table S4. The precipitation (`precip`) and relative humidity (`rh`) covariates are running means as described in main text. Other covariates include percent use of insecticide treated nets (`itn`) and human population density censored above 2,000 inhabitants per square kilometre (`hpd`). Covariates are scaled by minimum and maximum of observed values. Model estimates are shown with standard errors in parentheses.

| | |
|---|---|
| (Intercept) | $1.53^{***}$ |
| | (0.20) |
| light | $-1.05^{***}$ |
| | (0.09) |
| exit | $-1.69^{***}$ |
| | (0.11) |
| PSC | $-0.43^{**}$ |
| | (0.13) |
| animal | $-0.08$ |
| | (0.14) |
| pit | $-2.06^{***}$ |
| | (0.14) |
| outdoor | $-0.20$ |
| | (0.12) |
| precip | $6.04^{***}$ |
| | (0.47) |
| $\text{precip}^2$ | $-2.57^{**}$ |
| | (0.81) |
| rh | $1.45^{***}$ |
| | (0.23) |
| hpd | 0.20 |
| | (0.50) |
| $\text{hpd}^2$ | $-6.63^{***}$ |
| | (0.24) |
| itn | $-2.39^{***}$ |
| | (0.08) |
| precip:hpd | $-5.40^{***}$ |
| | (0.54) |
| rh:hpd | $8.19^{***}$ |
| | (0.65) |
| AIC | 59071.72 |
| BIC | 59185.59 |
| Log Likelihood | $-29519.86$ |
| Deviance | 9644.30 |
| Num. obs. | 9105 |

$^{***}p < 0.001$; $^{**}p < 0.01$; $^{*}p < 0.05$

### Spatial Location of Vector Occurrence and Count Data

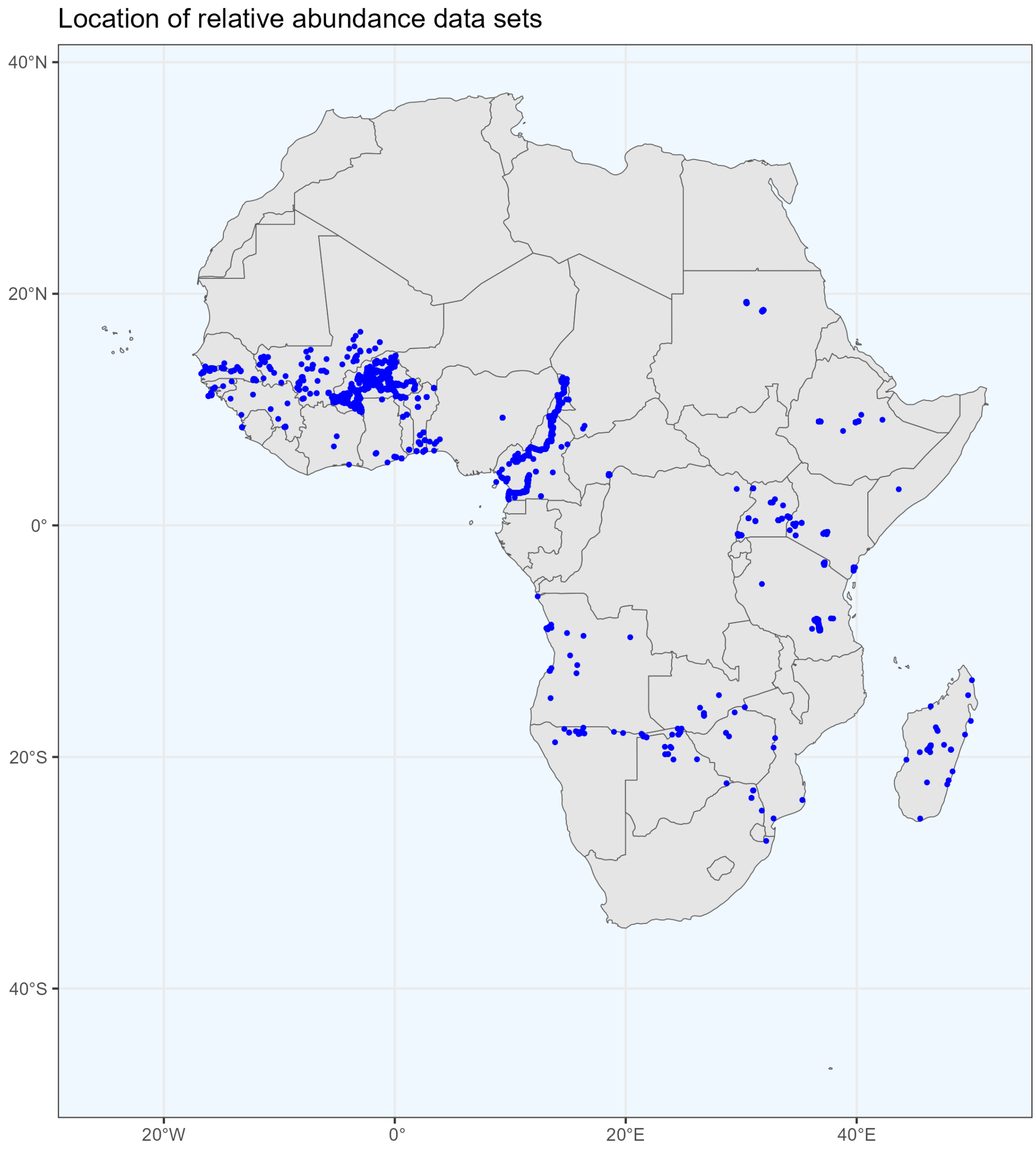


Figure S1: Spatial locations of data sources (Table S1) based on species occurrence data provided by `VectorBase.org` under CC BY 4.0 license. Made with Natural Earth. Vector map data were obtained from `naturalearthdata.com`.

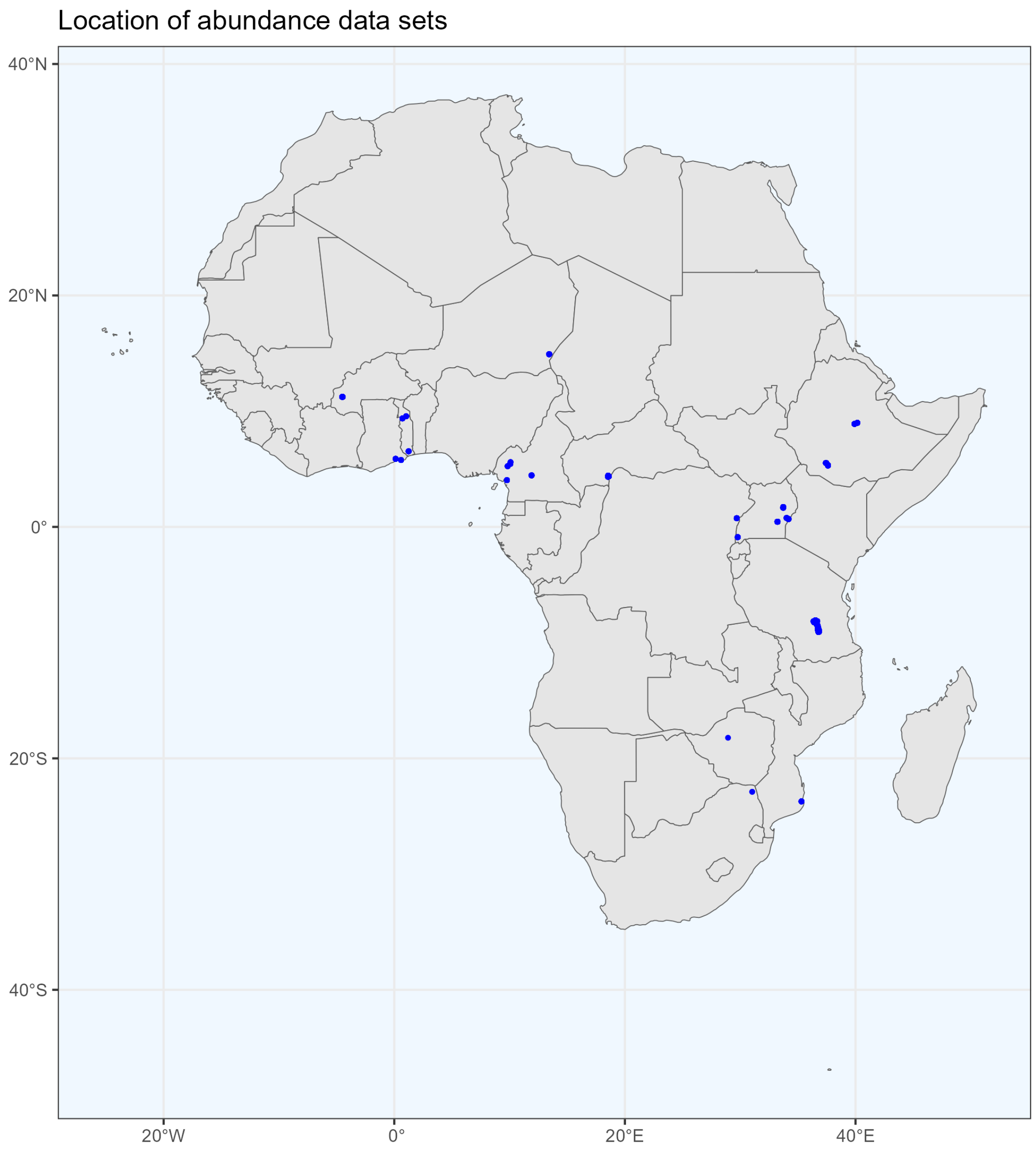


Figure S2: Spatial locations of data sources (Table S2) based on abundance data provided by `VectorBase.org` under CC BY 4.0 license. Made with Natural Earth. Vector map data were obtained from `naturalearthdata.com`.

### Covariate Visualisations

#### Meteorological Covariates

Meteorological covariates for precipitation, relative humidity and temperature entered into the models as daily means. For visualisation, annual averages are shown in Figures S3, S4 and S5. Quarterly averages are shown in Figures S6, S7 and S8. Annual anomalies are shown for four different years in Figures S9, S10 and S11.

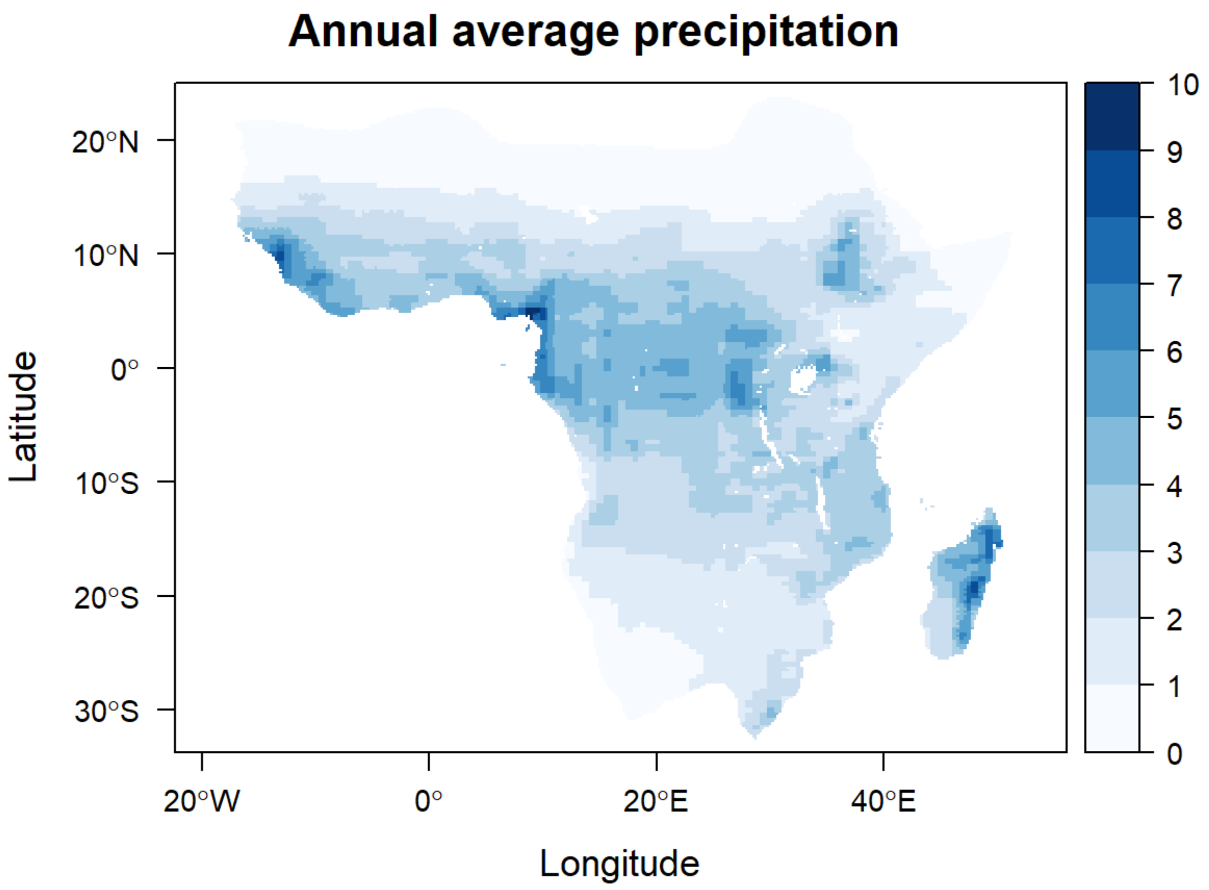


Figure S3: Annual average MERRA-2 daily precipitation (mm/day, corrected) over years 2002–2020 in the potential range of *An. gambiae s.l.* (1) as provided by the Malaria Atlas Project (`https://malariaatlas.org/`) under the terms of a Creative Commons Attribution 4.0 International License available at `https://creativecommons.org/licenses/by/4.0/`. These data were obtained from the NASA Langley Research Center (LaRC) POWER Project funded through the NASA Earth Science/Applied Science Program.

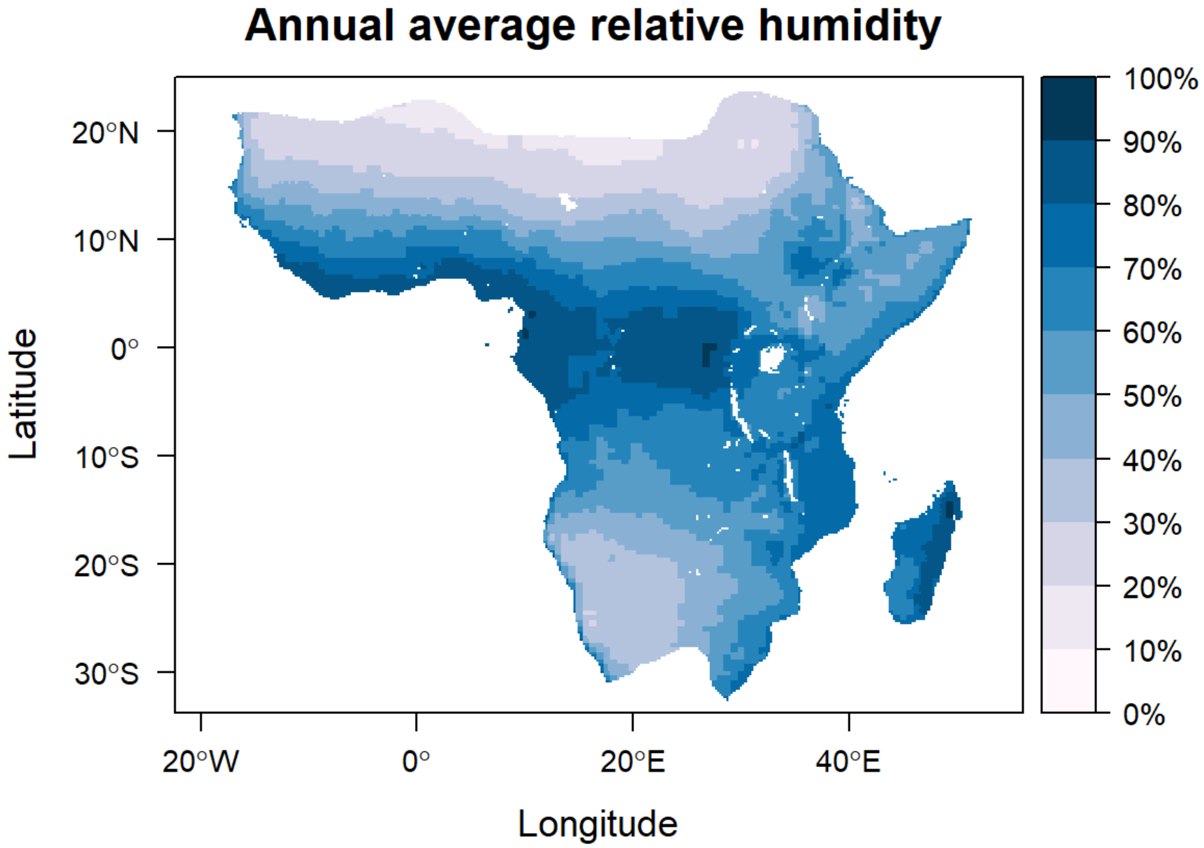


Figure S4: Annual average MERRA-2 daily relative humidity at 2 metres (%) over years 2002–2020 in the potential range of *An. gambiae s.l.* (1) as provided by the Malaria Atlas Project (`https://malariaatlas.org/`) under the terms of a Creative Commons Attribution 4.0 International License available at `https://creativecommons.org/licenses/by/4.0/`. These data were obtained from the NASA Langley Research Center (LaRC) POWER Project funded through the NASA Earth Science/Applied Science Program.

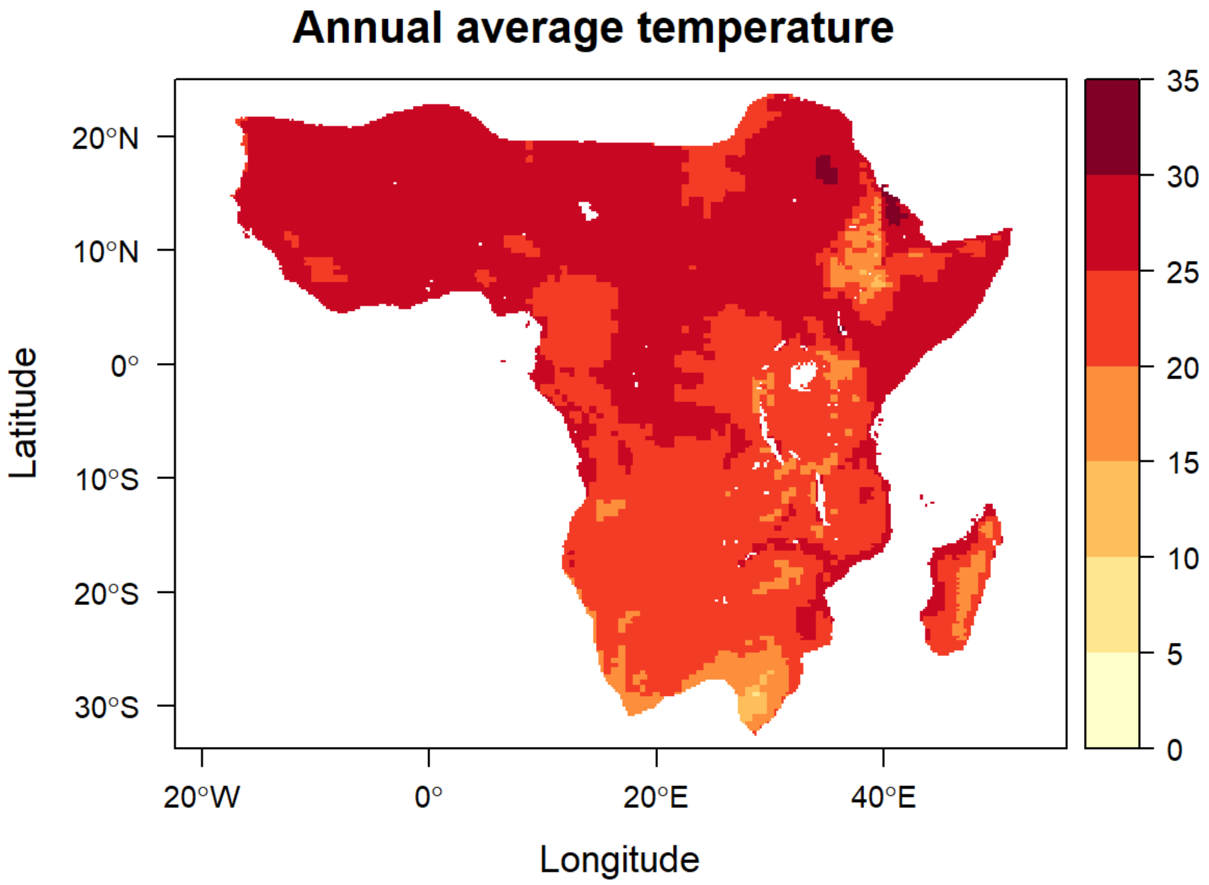


Figure S5: Annual average MERRA-2 daily temperature at 2 metres (degrees C) over years 2002–2020 in the potential range of *An. gambiae s.l.* (1) as provided by the Malaria Atlas Project (`https://malariaatlas.org/`) under the terms of a Creative Commons Attribution 4.0 International License available at `https://creativecommons.org/licenses/by/4.0/`. These data were obtained from the NASA Langley Research Center (LaRC) POWER Project funded through the NASA Earth Science/Applied Science Program.

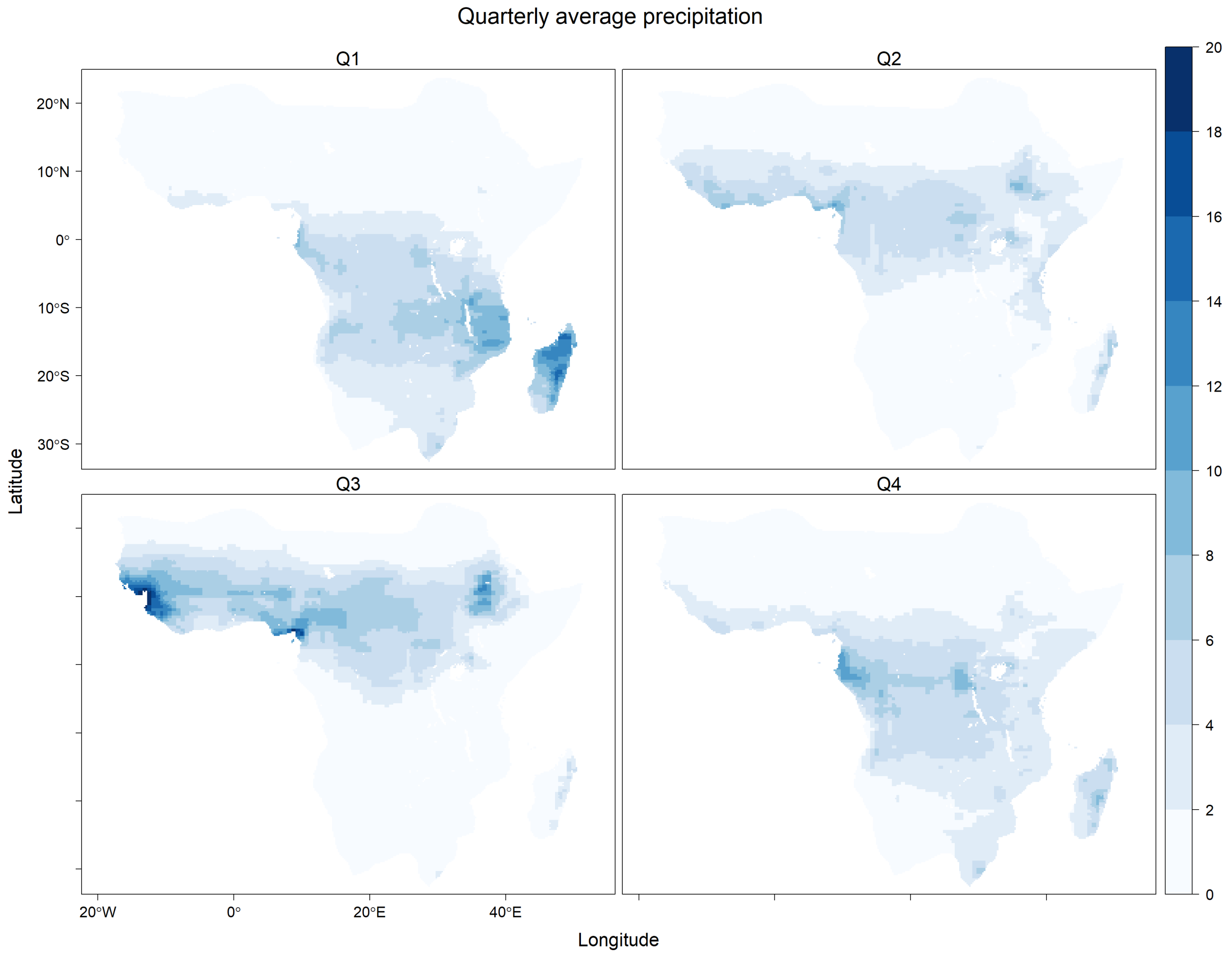


Figure S6: Average MERRA-2 daily precipitation (mm/day, corrected) by quarter over years 2002–2020 in the potential range of *An. gambiae s.l.* (1) as provided by the Malaria Atlas Project (`https://malariaatlas.org/`) under the terms of a Creative Commons Attribution 4.0 International License available at `https://creativecommons.org/licenses/by/4.0/`. These data were obtained from the NASA Langley Research Center (LaRC) POWER Project funded through the NASA Earth Science/Applied Science Program.

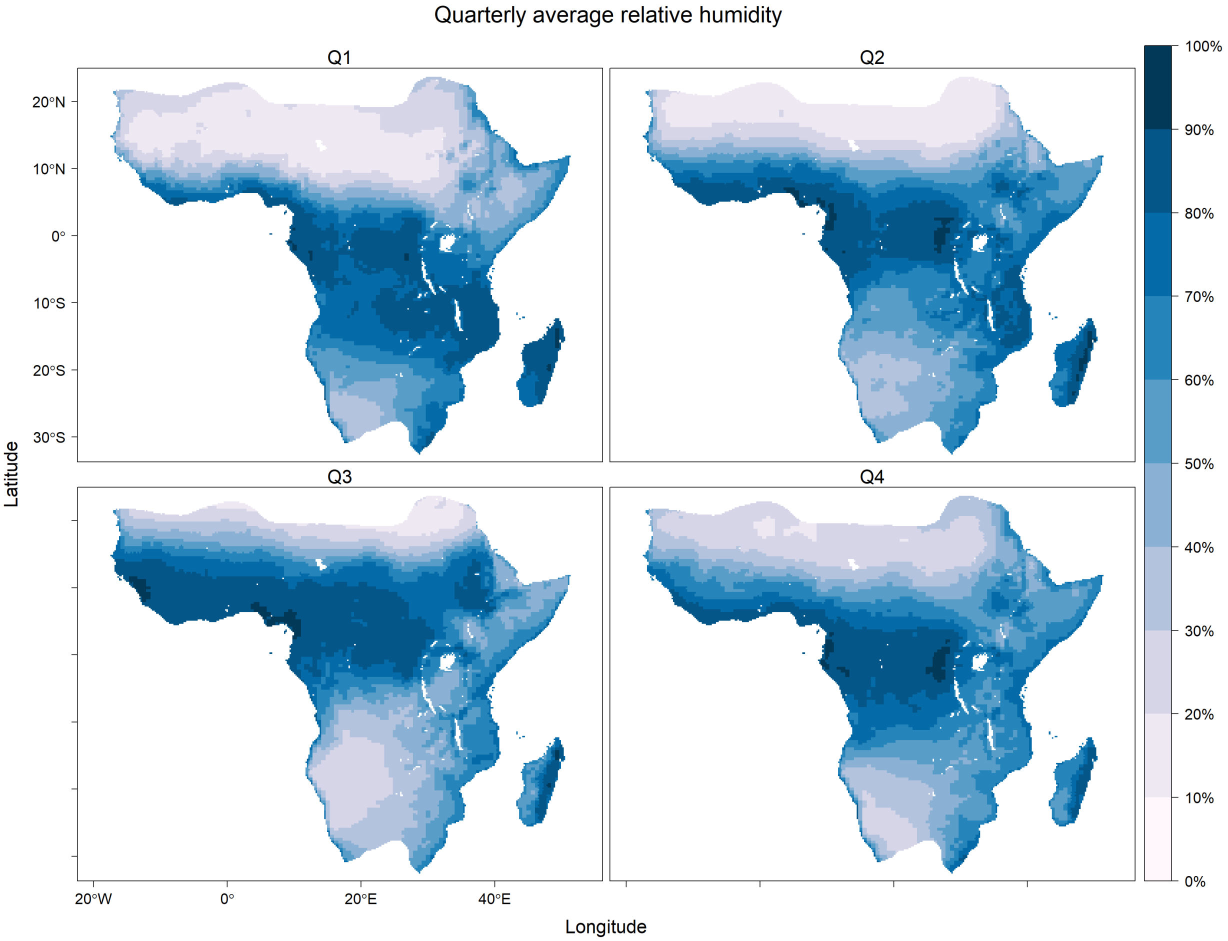


Figure S7: Average MERRA-2 daily relative humidity at 2 metres (%) by quarter over years 2002–2020 in the potential range of *An. gambiae s.l.* (1) as provided by the Malaria Atlas Project (`https://malariaatlas.org/`) under the terms of a Creative Commons Attribution 4.0 International License available at `https://creativecommons.org/licenses/by/4.0/`. These data were obtained from the NASA Langley Research Center (LaRC) POWER Project funded through the NASA Earth Science/Applied Science Program.

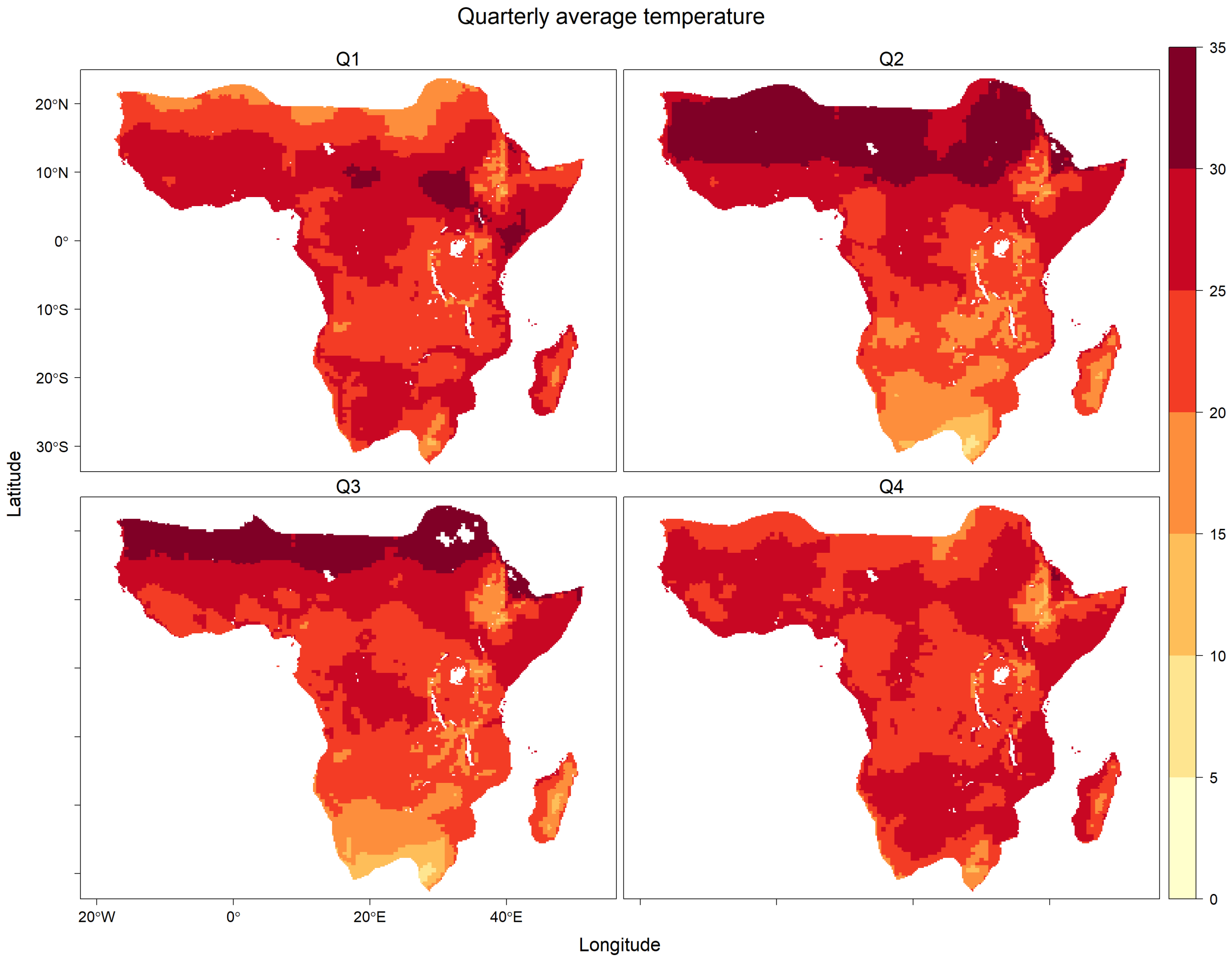


Figure S8: Average MERRA-2 daily temperature at 2 metres (degrees C) by quarter over years 2002–2020 in the potential range of *An. gambiae s.l.* (1) as provided by the Malaria Atlas Project (`https://malariaatlas.org/`) under the terms of a Creative Commons Attribution 4.0 International License available at `https://creativecommons.org/licenses/by/4.0/`. These data were obtained from the NASA Langley Research Center (LaRC) POWER Project funded through the NASA Earth Science/Applied Science Program.

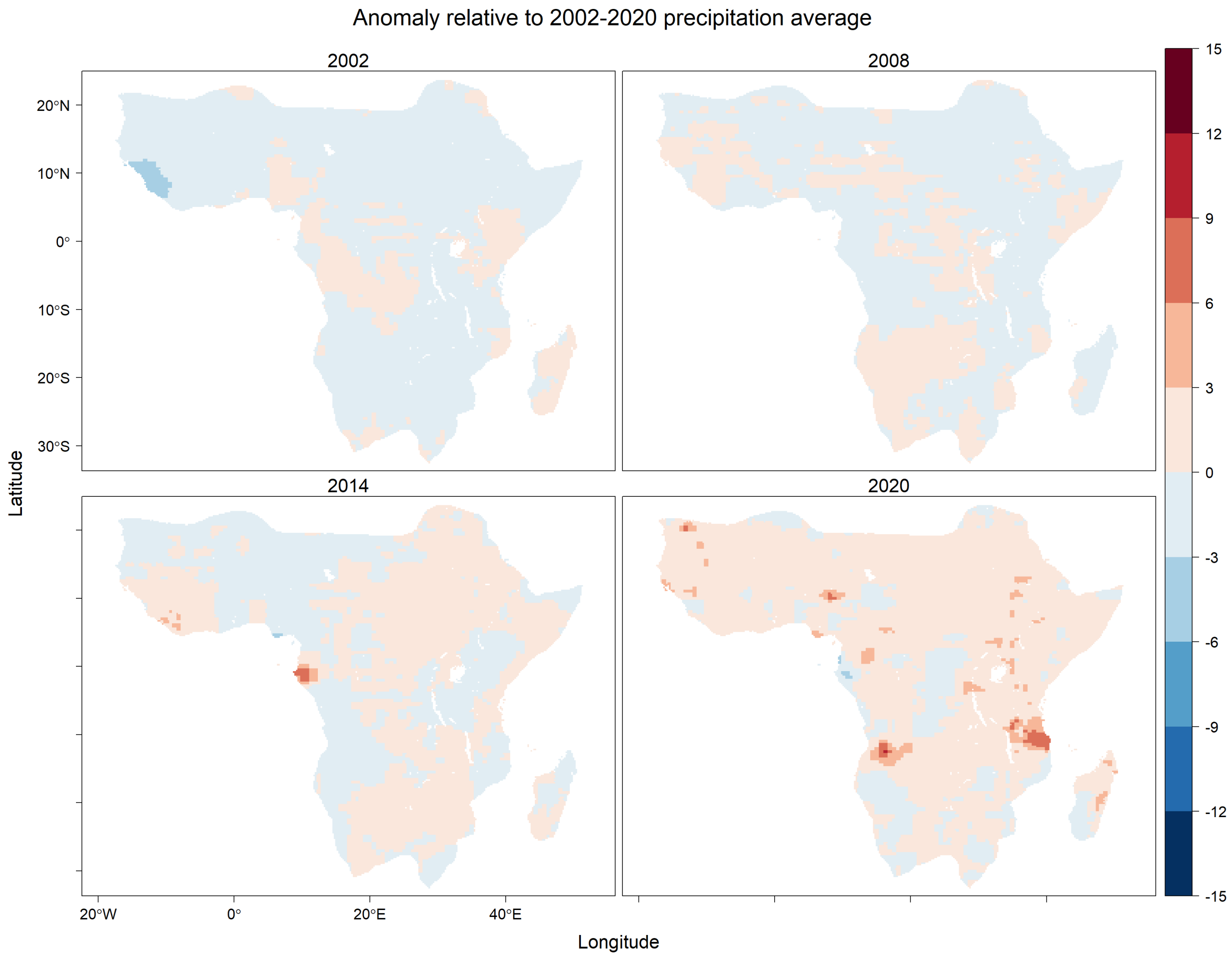


Figure S9: Annual average differences of MERRA-2 daily precipitation (mm/day, corrected) with respect to average over years 2002–2020 in the potential range of *An. gambiae s.l.* (1) as provided by the Malaria Atlas Project (`https://malariaatlas.org/`) under the terms of a Creative Commons Attribution 4.0 International License available at `https://creativecommons.org/licenses/by/4.0/`. These data were obtained from the NASA Langley Research Center (LaRC) POWER Project funded through the NASA Earth Science/Applied Science Program

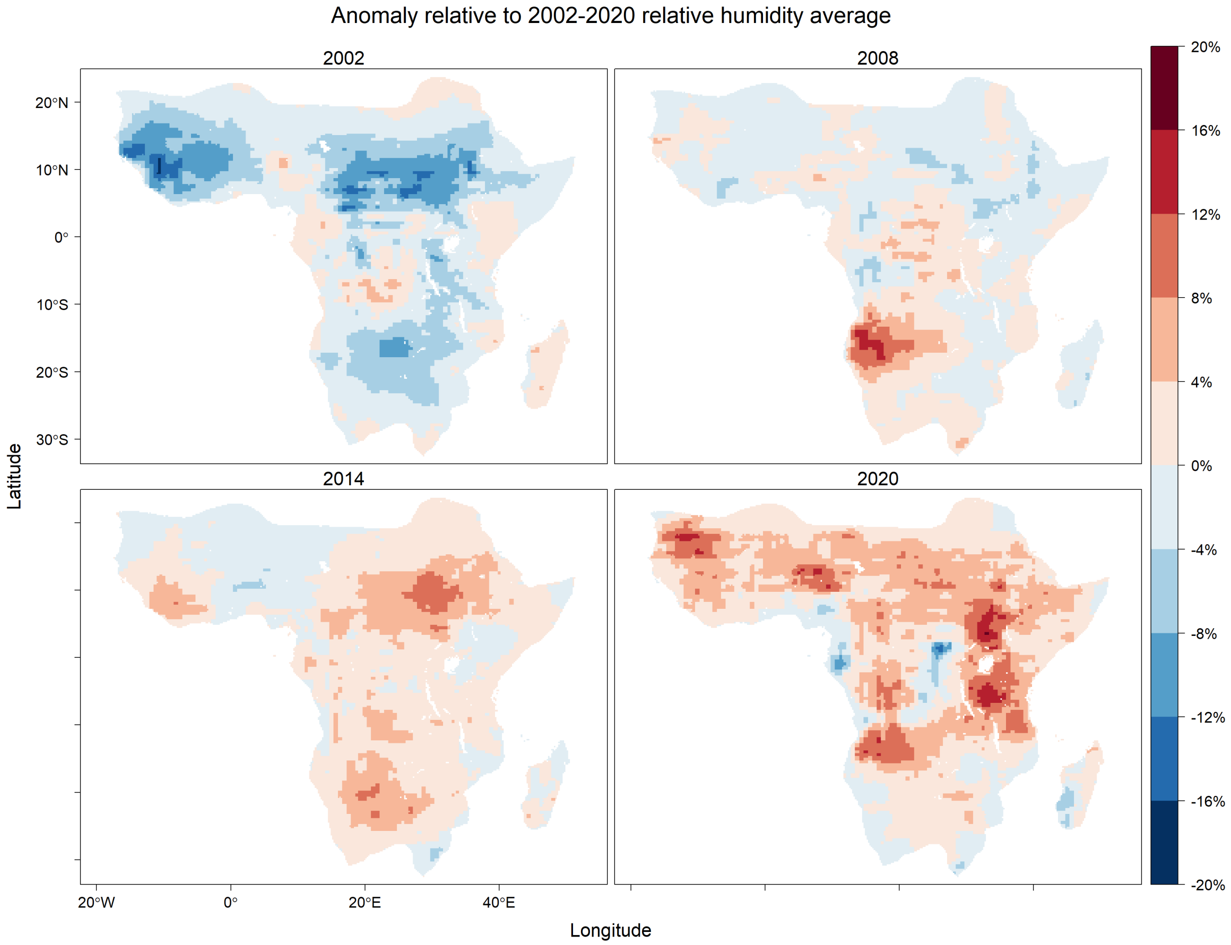


Figure S10: Annual average differences of MERRA-2 daily relative humidity at 2 metres (%) with respect to average over years 2002–2020 in the potential range of *An. gambiae s.l.* (1) as provided by the Malaria Atlas Project (`https://malariaatlas.org/`) under the terms of a Creative Commons Attribution 4.0 International License available at `https://creativecommons.org/licenses/by/4.0/`. These data were obtained from the NASA Langley Research Center (LaRC) POWER Project funded through the NASA Earth Science/Applied Science Program.

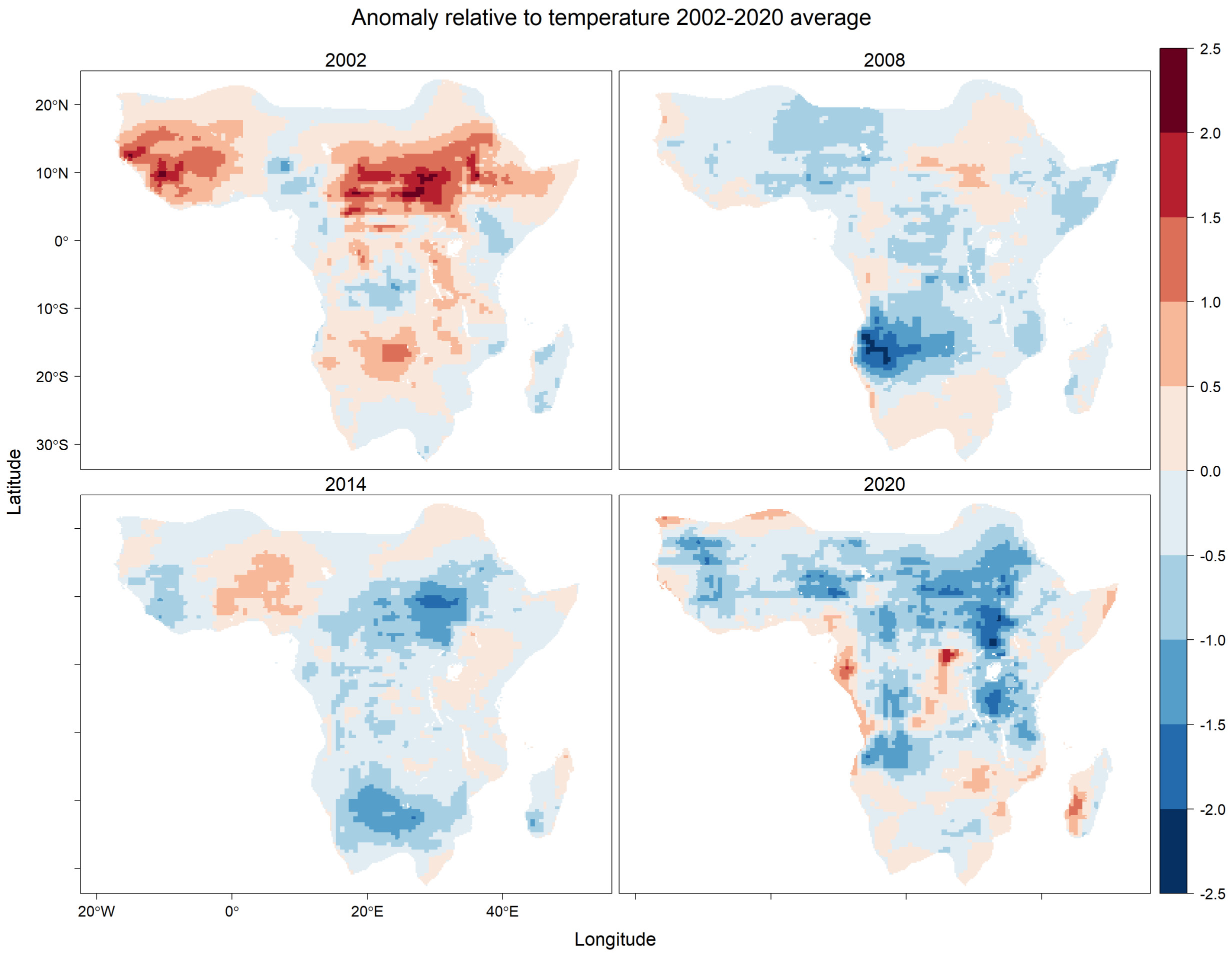


Figure S11: Annual average difference of MERRA-2 daily temperature at 2 metres (degrees C) with respect to average over years 2002–2020 in the potential range of *An. gambiae s.l.* (1) as provided by the Malaria Atlas Project (`https://malariaatlas.org/`) under the terms of a Creative Commons Attribution 4.0 International License available at `https://creativecommons.org/licenses/by/4.0/`. These data were obtained from the NASA Langley Research Center (LaRC) POWER Project funded through the NASA Earth Science/Applied Science Program.

#### Geographical Covariates

Visualisations are shown for the spatial covariates of distance to coastline (Figure S12), probability of permanent freshwater (Figure S14), elevation (Figure S13) and human population density (Figure S15).

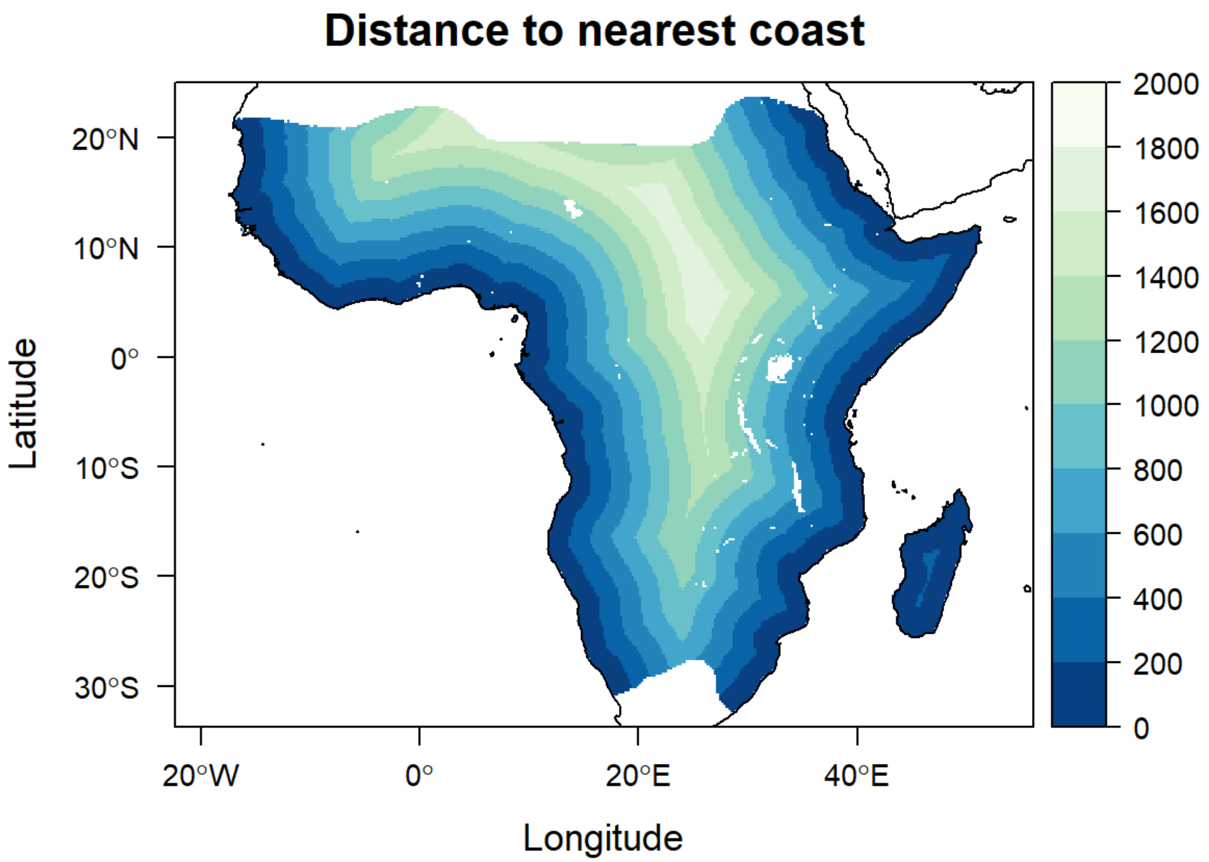


Figure S12: Distance to coastline (km) masked by potential range of *An. gambiae s.l.* (1) as provided by the Malaria Atlas Project (`https://malariaatlas.org/`) under the terms of a Creative Commons Attribution 4.0 International License available at `https://creativecommons.org/licenses/by/4.0/`. Made with Natural Earth. Free vector and raster map data were obtained from `naturalearthdata.com`. The naturalearth data are licensed PD CC, see `https://creativecommons.org/publicdomain/`.

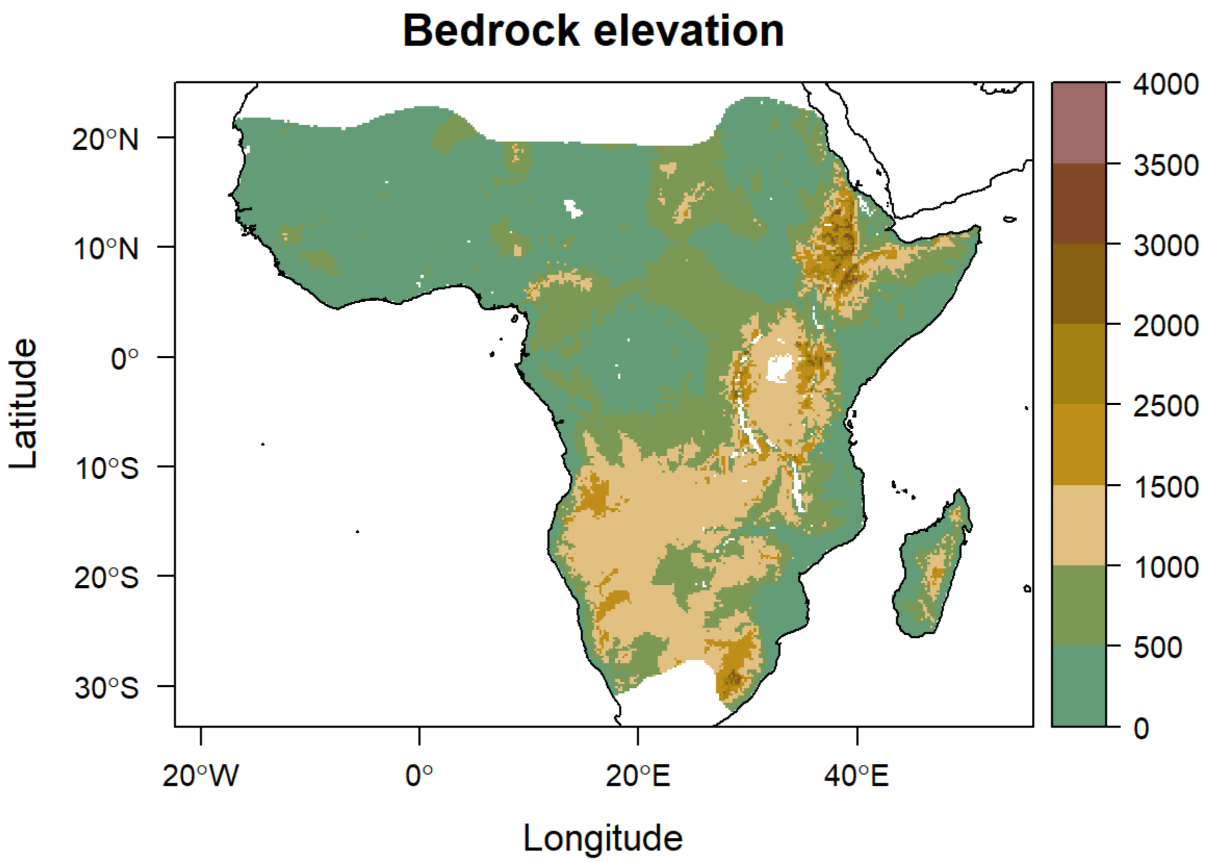


Figure S13: Bedrock elevation (metres) masked by potential range of *An. gambiae s.l.* (1) as provided by the Malaria Atlas Project (`https://malariaatlas.org/`) under the terms of a Creative Commons Attribution 4.0 International License available at `https://creativecommons.org/licenses/by/4.0/`. Elevation data from NOAA NCEI (4).

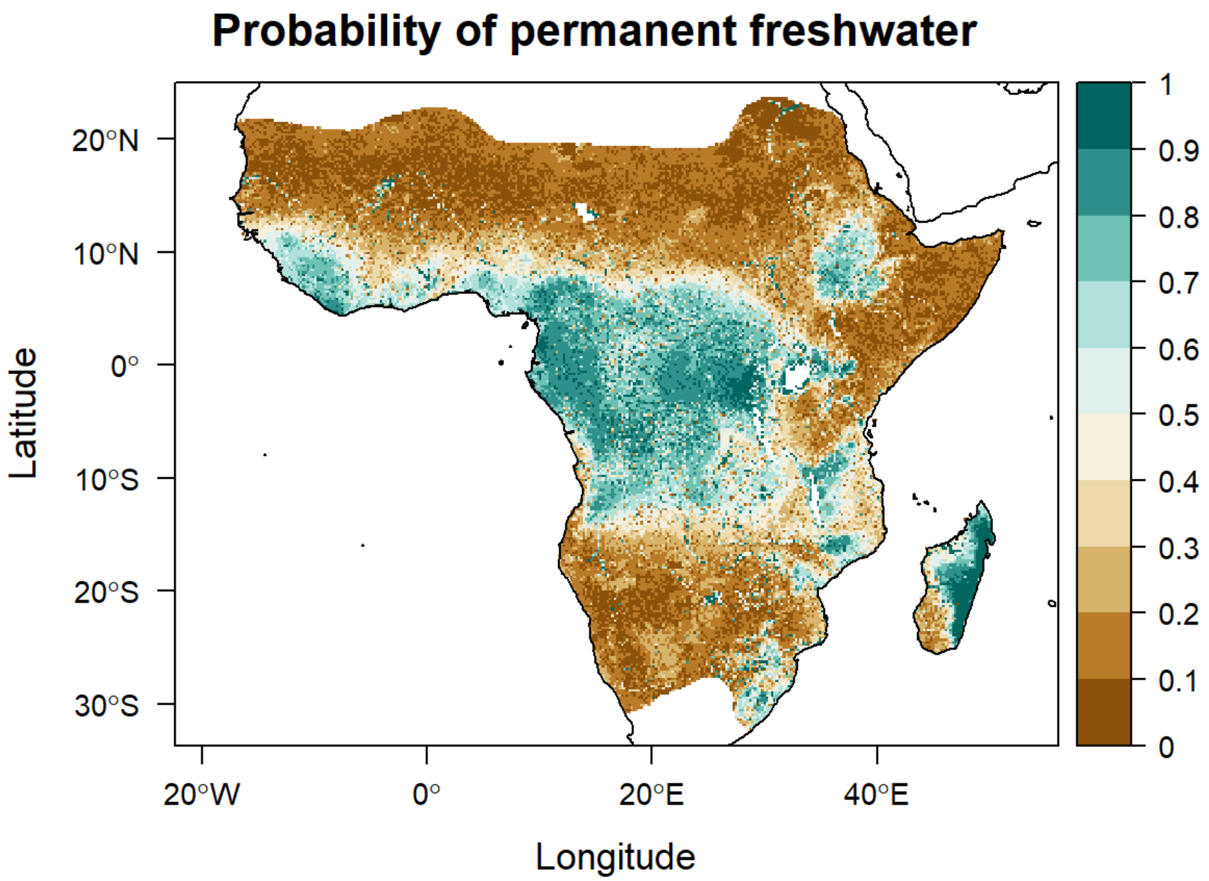


Figure S14: Probability of permanent freshwater derived from lakes data (2), rivers and streams data (3), and masked by potential range of *An. gambiae s.l.* (1) as provided by the Malaria Atlas Project (`https://malariaatlas.org/`). Data provided under the terms of a Creative Commons Attribution 4.0 International License available at `https://creativecommons.org/licenses/by/4.0/`.

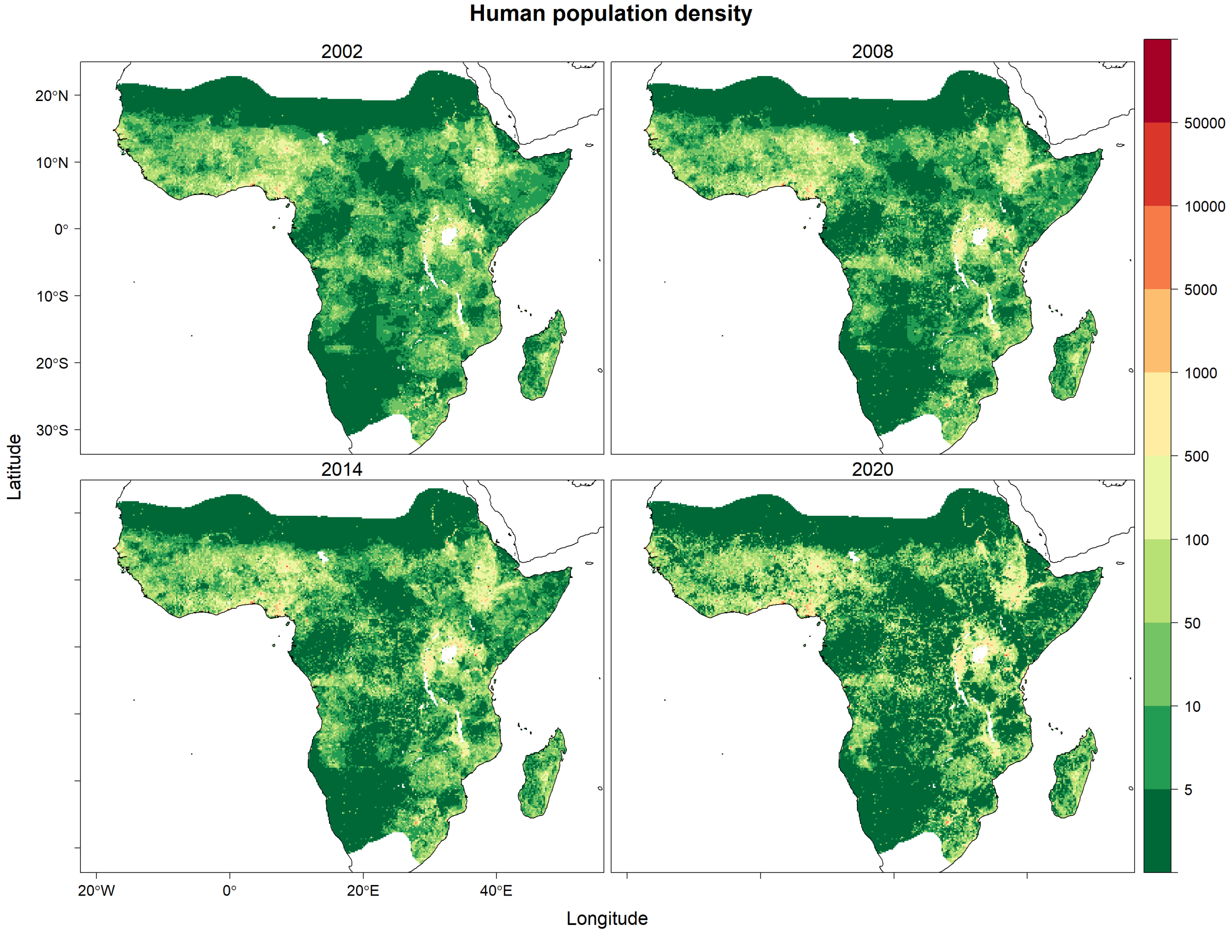


Figure S15: Average human population density per square kilometre in four selected years, masked by potential range of *An. gambiae s.l.* (1) as provided by the Malaria Atlas Project (`https://malariaatlas.org/`) under the terms of a Creative Commons Attribution 4.0 International License available at `https://creativecommons.org/licenses/by/4.0/`. Human population data from LANDSCAN (5; 6; 7; 8) provided under CC BY 4.0 license.

#### Vector Intervention Covariates

Annual vector intervention data available for the prediction period 2002–2020 are visualised for four different years in Figures S16 and S17.

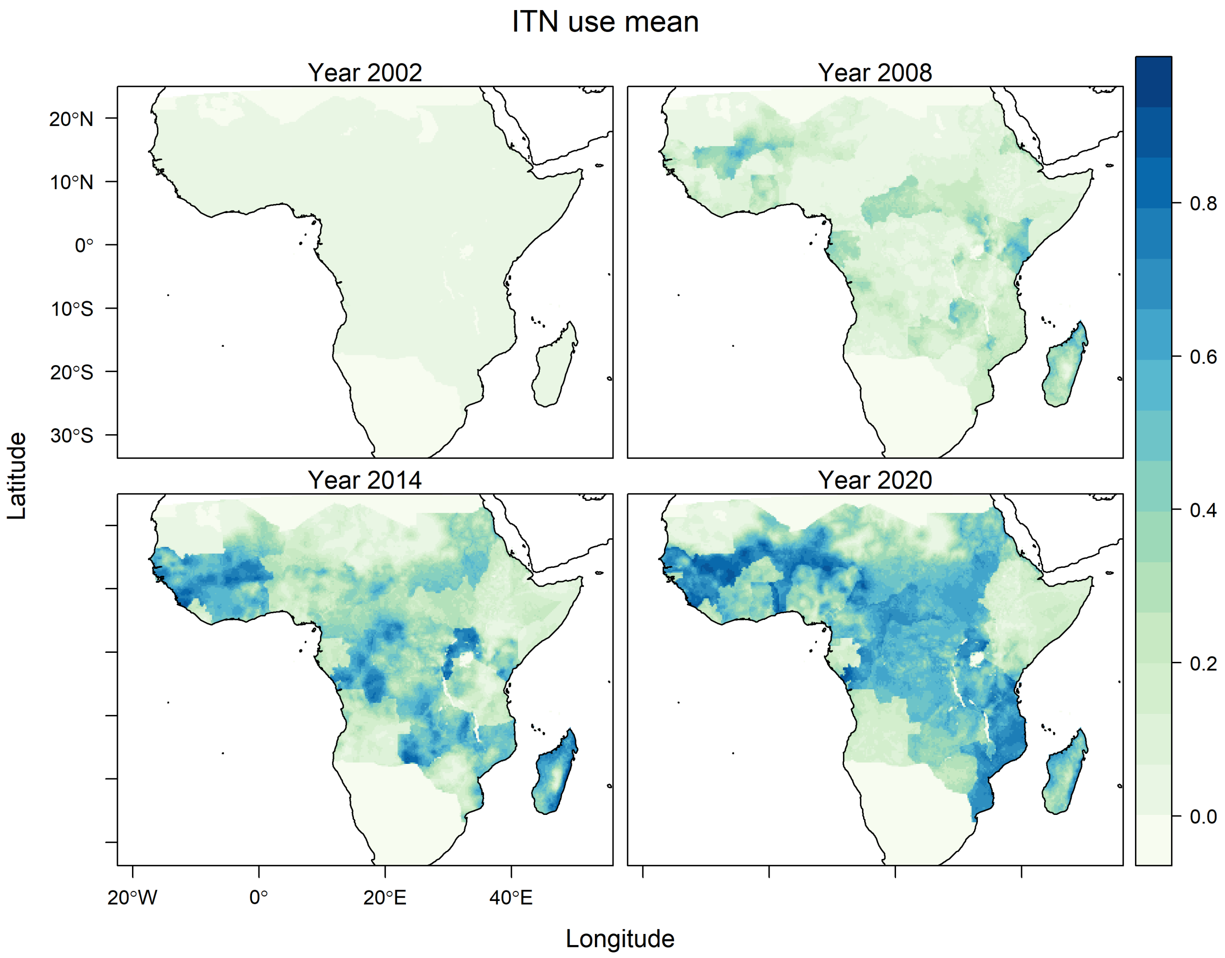


Figure S16: Annual proportion of insecticide treated net (ITN) use as provided by the Malaria Atlas Project (`https://malariaatlas.org/`) under the terms of a Creative Commons Attribution 4.0 International License available at `https://creativecommons.org/licenses/by/4.0/`.

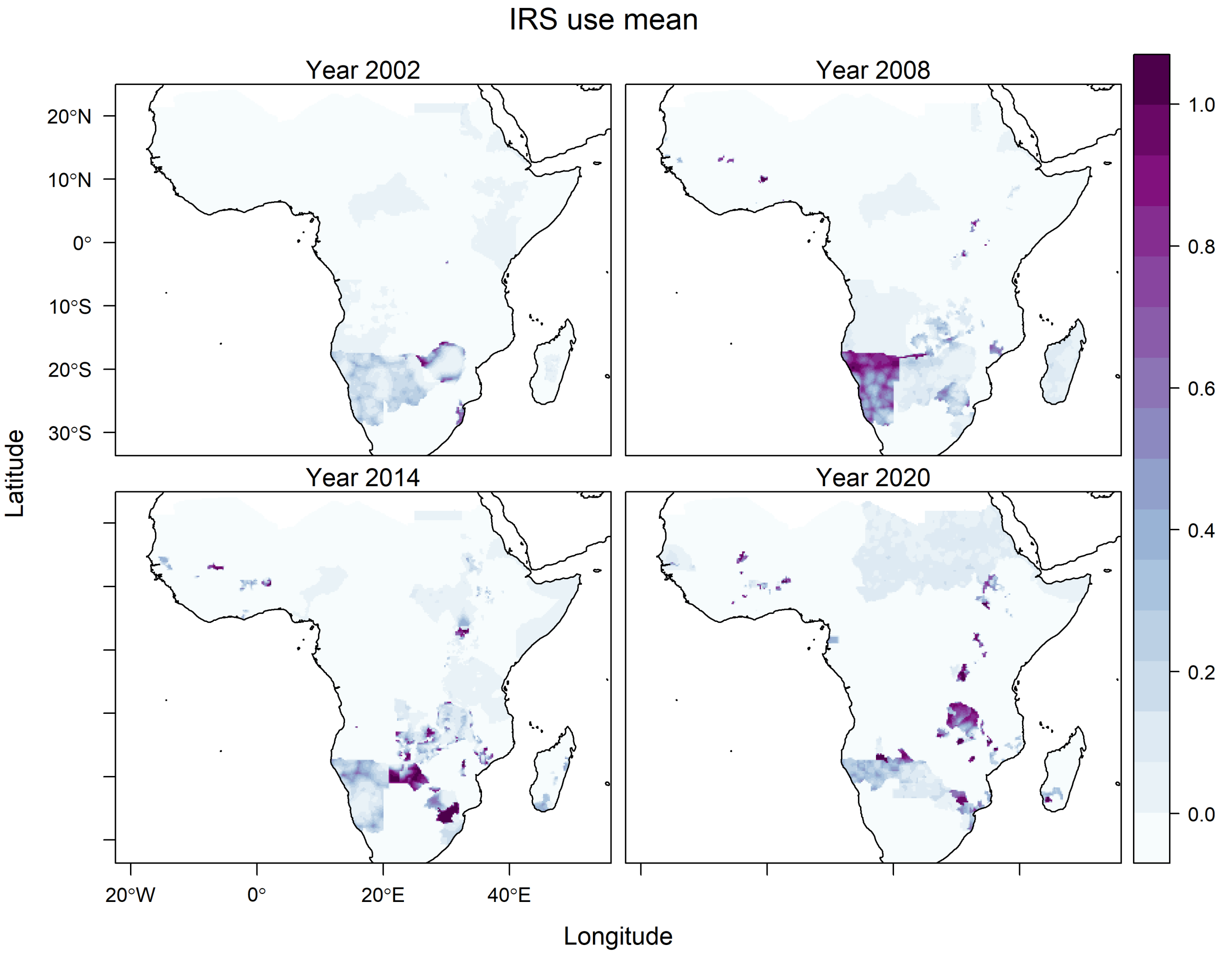


Figure S17: Annual proportion of houses covered by indoor residual spraying (IRS) as provided by the Malaria Atlas Project (`https://malariaatlas.org/`) under the terms of a Creative Commons Attribution 4.0 International License available at `https://creativecommons.org/licenses/by/4.0/`.

### Annual Predictions

Figures S18, S19 and S20 show annual anomalies in predicted equilibrium abundance for four different years during the prediction period 2002–2020.

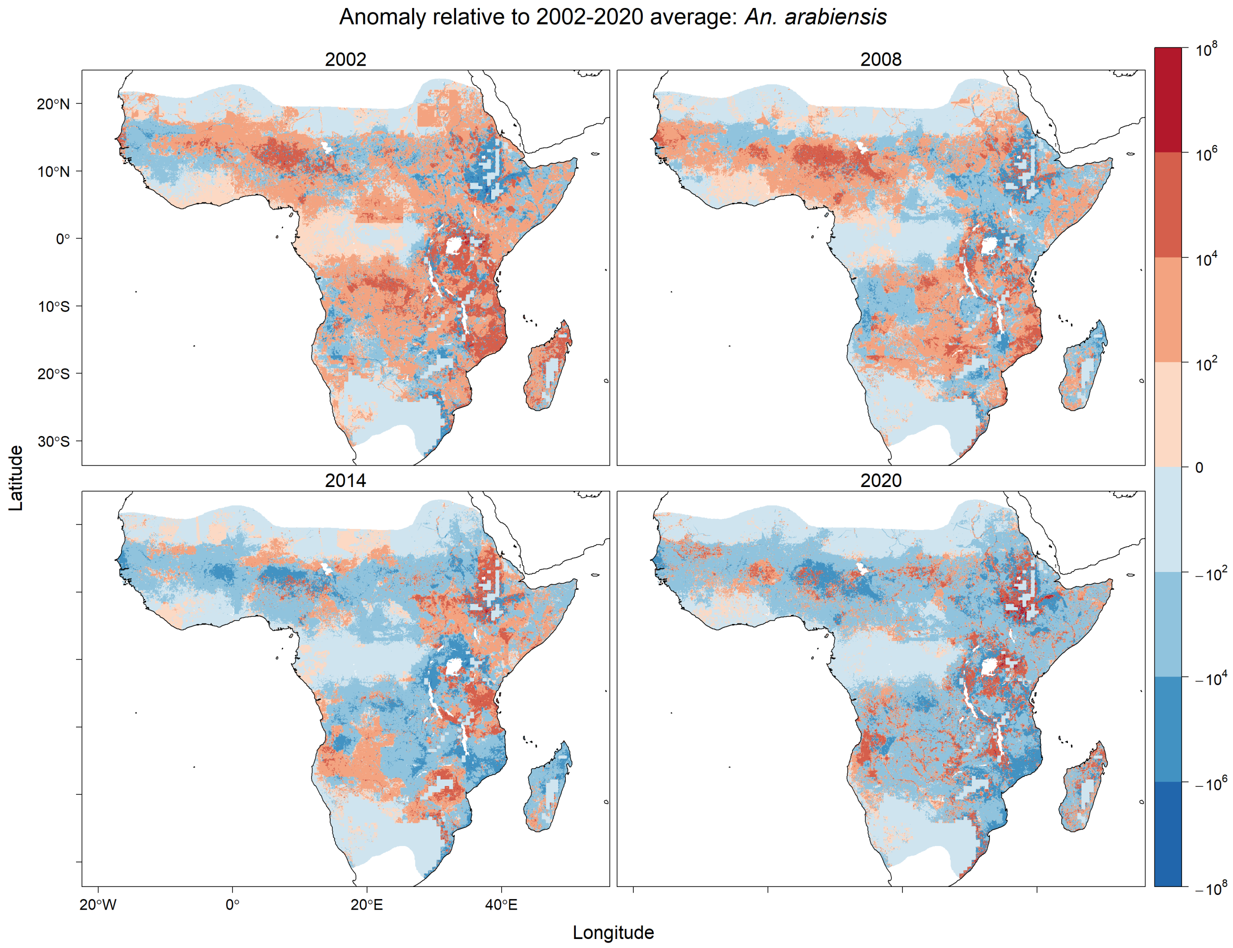


Figure S18: Annual average anomalies of total potential equilibrium vector abundance of *An. arabiensis* relative to its predicted average per 25 square kilometres over years 2002–2020. Predictions are masked to the potential range of *An. gambiae s.l.* (1) provided by the Malaria Atlas Project (`https://malariaatlas.org/`) under the terms of a Creative Commons Attribution 4.0 International License available at `https://creativecommons.org/licenses/by/4.0/`.

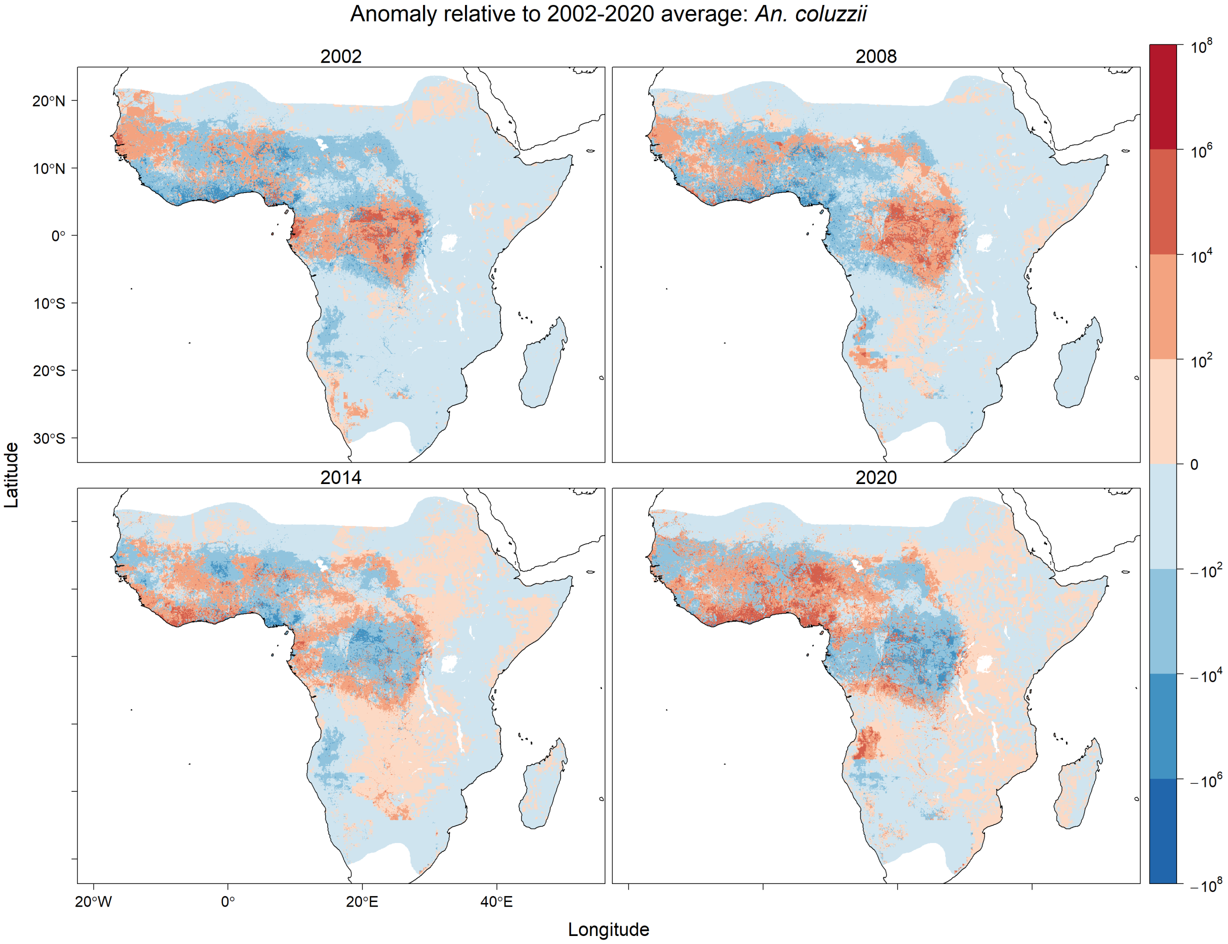


Figure S19: Annual average anomalies of total potential equilibrium vector abundance of *An. coluzzii* relative to its predicted average per 25 square kilometres over years 2002–2020. Predictions are masked to the potential range of *An. gambiae s.l.* (1) provided by the Malaria Atlas Project (`https://malariaatlas.org/`) under the terms of a Creative Commons Attribution 4.0 International License available at `https://creativecommons.org/licenses/by/4.0/`.

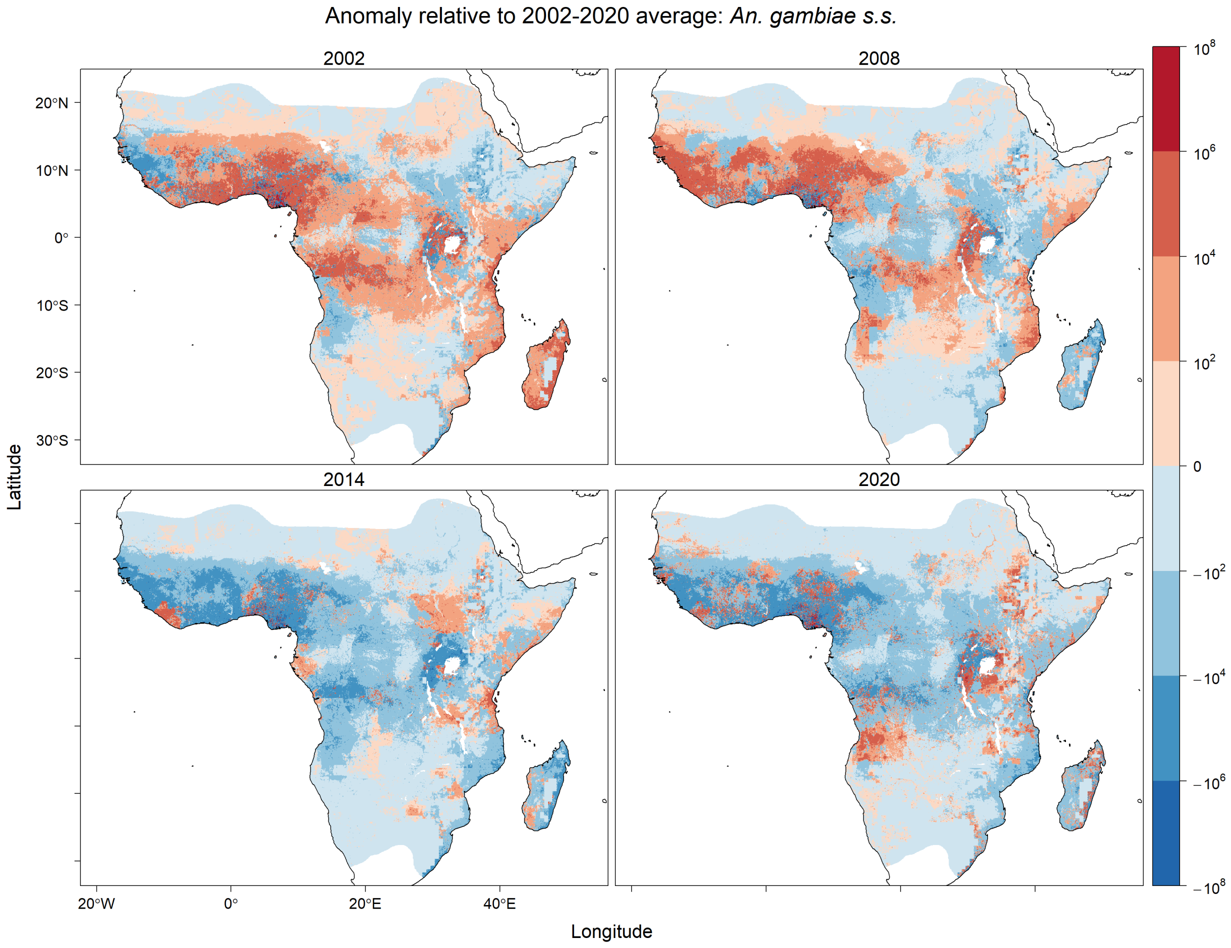


Figure S20: Annual average anomalies of total potential equilibrium vector abundance of *An. gambiae s.s.* relative to its predicted average per 25 square kilometres over years 2002–2020. Predictions are masked to the potential range of *An. gambiae s.l.* (1) provided by the Malaria Atlas Project (`https://malariaatlas.org/`) under the terms of a Creative Commons Attribution 4.0 International License available at `https://creativecommons.org/licenses/by/4.0/`.

#### Predictions from Generalised Linear Models

The multinomial logit GLM predictions of species composition are shown in Figure S21 averaged over the prediction period 2002–2020 in each cell. The associated average prediction variance is shown in Figure S22. The per capita predictions of vector abundance of *An. gambiae* sensu lato are shown in Figure S24a averaged over the prediction period in each cell. The associated average prediction variance is shown in Figure S24b.

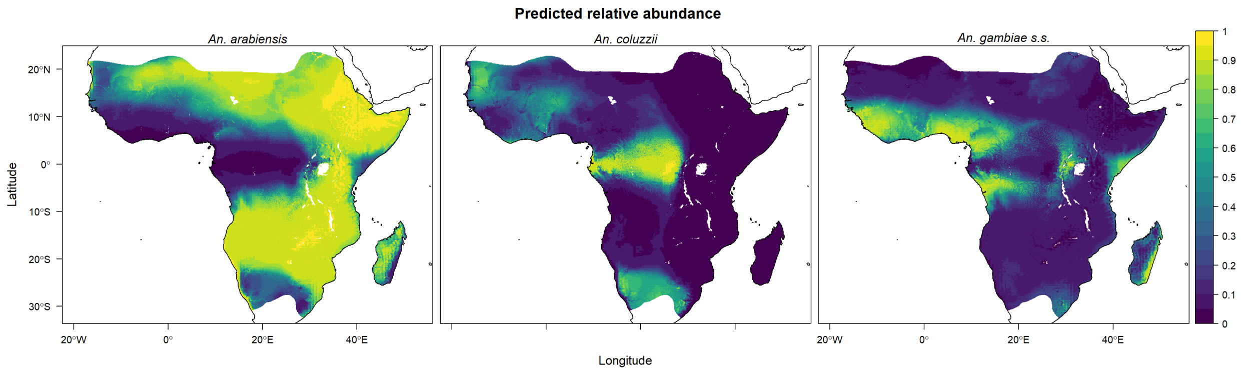


Figure S21: Mean predicted relative abundance of *An. arabiensis*, *An. coluzzii* and *An. gambiae* s.s. over years 2002–2020 from the multinomial logit GLM. Note that the predictions shown in this map do not account for temporally varying abundance of *An. gambiae s.l.* Predictions are masked to the potential range of *An. gambiae s.l.* (1) provided by the Malaria Atlas Project (`https://malariaatlas.org/`) under the terms of a Creative Commons Attribution 4.0 International License available at `https://creativecommons.org/licenses/by/4.0/`.

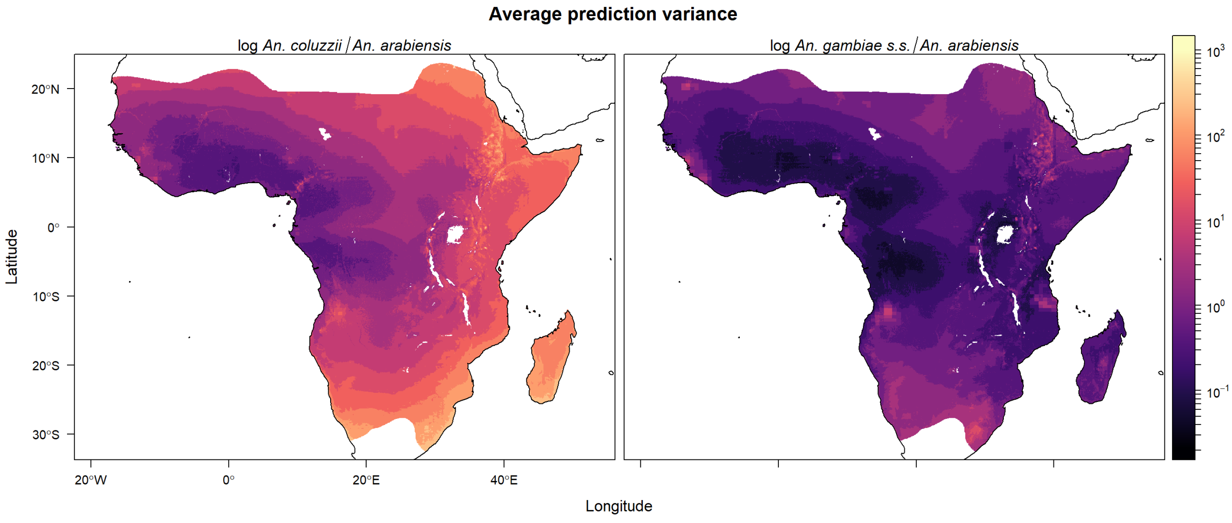


Figure S22: Average prediction variance of linear predictor for relative abundance of *An. gambiae* sensu lato over years 2002–2020 in the multinomial GLM. Predictions are masked to the potential range of *An. gambiae s.l.* (1) provided by the Malaria Atlas Project (`https://malariaatlas.org/`) under the terms of a Creative Commons Attribution 4.0 International License available at `https://creativecommons.org/licenses/by/4.0/`. Note logarithmic scale.

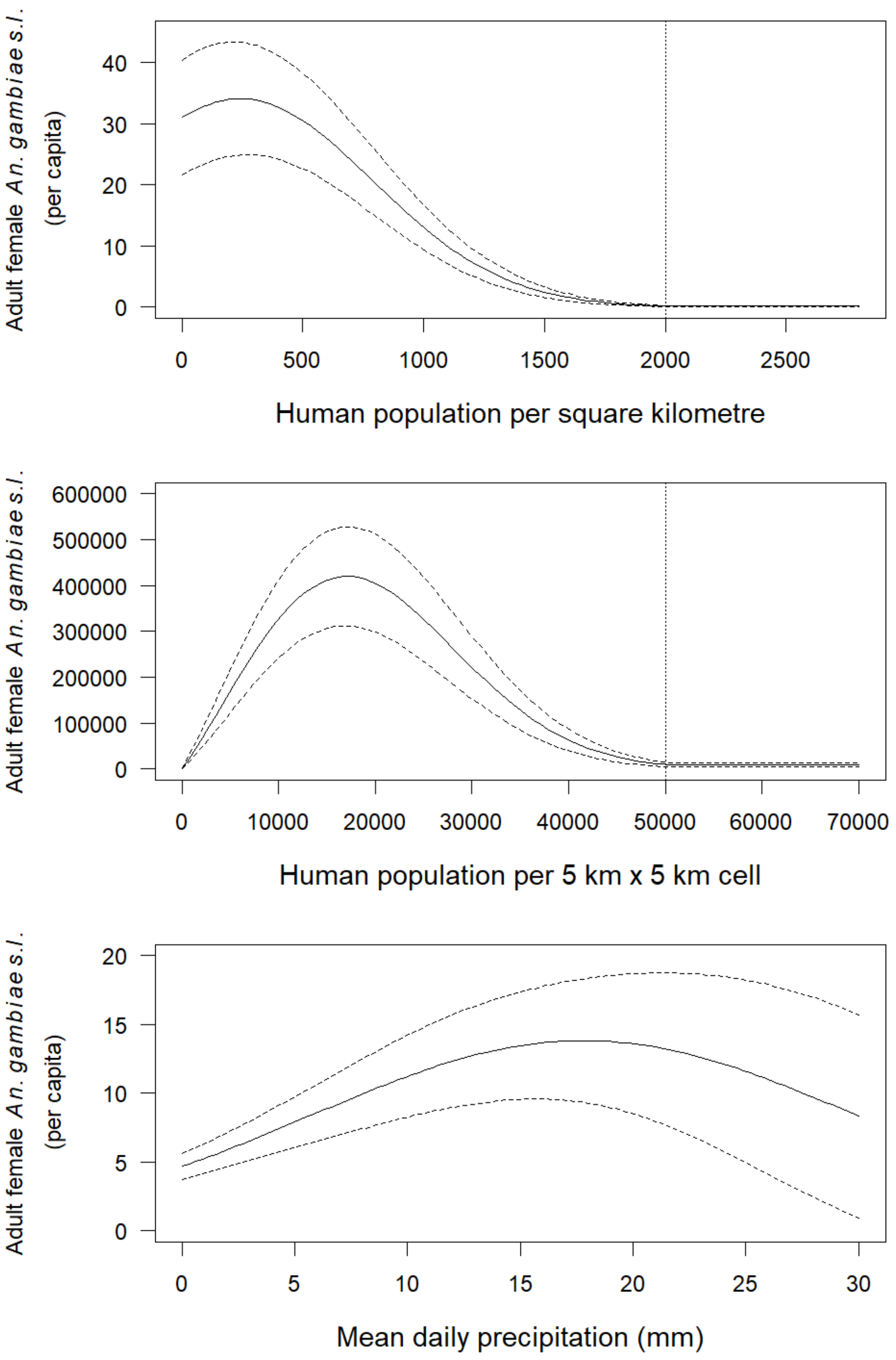


Figure S23: Prediction of vector abundance of *An. gambiae* sensu lato: Per capita adult female vector abundance with varying human population density (top), total female vector abundance with varying total human population per 5 km × 5 km grid cell (middle), and per capita adult female vector abundance with varying precipitation (bottom). Mean predictions plus or minus one standard error are shown. Vertical dotted lines depict the threshold for urban populations. Non-varying covariates in each plot are held constant at the midpoints of the within-sample observed ranges: 13.7 mm mean precipitation per day (for varying human population), 1000 humans per square km or equivalently 25,000 humans per 5 km × 5 km grid cell (for varying precipitation), 50% mean relative humidity, and 43% insecticide treated net coverage.

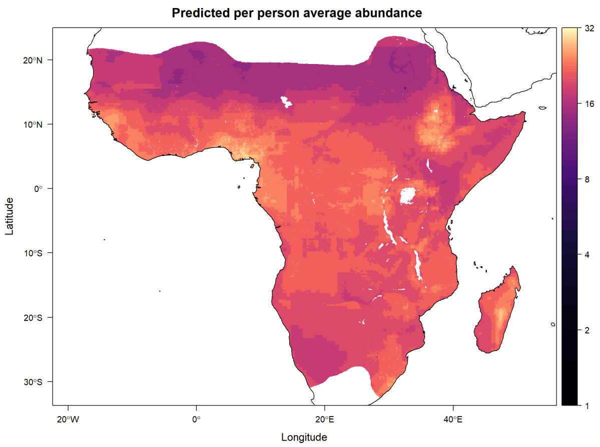


(a) Mean prediction total per person vector abundance of *An. gambiae* sensu lato over years 2002–2020.

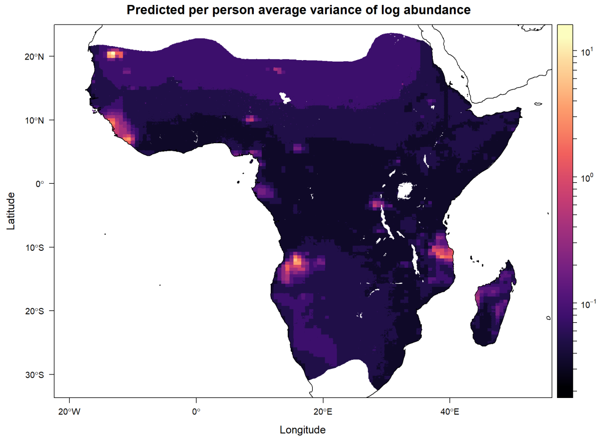


(b) Variance of linear predictor for total per person vector abundance of *An. gambiae* sensu lato over years 2002–2020.

Figure S24: Per capita predictions of vector abundance of *An. gambiae* sensu lato. Predictions are masked to the potential range of *An. gambiae s.l.* (1) provided by the Malaria Atlas Project (`https://malariaatlas.org/`) under the terms of a Creative Commons Attribution 4.0 International License available at `https://creativecommons.org/licenses/by/4.0/`. Note logarithmic scale.
